# Spike protein mutational landscape in India: Could Muller’s ratchet be a future game-changer for COVID-19?

**DOI:** 10.1101/2020.08.18.255570

**Authors:** Rachana Banerjee, Kausik Basak, Anamika Ghosh, Vyshakh Rajachandran, Kamakshi Sureka, Debabani Ganguly, Sujay Chattopadhyay

## Abstract

The dire need of effective preventive measures and treatment approaches against SARS-CoV-2 virus, causing COVID-19 pandemic, calls for an in-depth understanding of its evolutionary dynamics with attention to specific geographic locations, since lockdown and social distancing to prevent the virus spread could lead to distinct localized dynamics of virus evolution within and between countries owing to different environmental and host-specific selection pressures. To decipher any correlation between SARS-CoV-2 evolution and its epidemiology in India, we studied the mutational diversity of spike glycoprotein, the key player for the attachment, fusion and entry of virus to the host cell. For this, we analyzed the sequences of 630 Indian isolates as available in GISAID database till June 07, 2020, and detected the spike protein variants to emerge from two major ancestors – Wuhan-Hu-1/2019 and its D614G variant. Average stability of the docked spike protein – host receptor (S-R) complexes for these variants correlated strongly (R^2^=0.96) with the fatality rates across Indian states. However, while more than half of the variants were found unique to India, 67% of all variants showed lower stability of S-R complex than the respective ancestral variants, indicating a possible fitness loss in recently emerged variants, despite a continuous increase in mutation rate. These results conform to the sharply declining fatality rate countrywide (>7-fold during April 11 – June 28, 2020). Altogether, while we propose the potential of S-R complex stability to track disease severity, we urge an immediate need to explore if SARS-CoV-2 is approaching mutational meltdown in India.

**Author summary:** Epidemiological features are intricately linked to evolutionary diversity of rapidly evolving pathogens, and SARS-CoV-2 is no exception. Our work suggests the potential of average stability of complexes formed by the circulating spike mutational variants and the human host receptor to track the severity of SARS-CoV-2 infection in a given region. In India, the stability of these complexes for recent variants tend to decrease relative to their ancestral ones, following countrywide declining fatality rate, in contrast to an increasing mutation rate. We hypothesize such a scenario as nascent footprints of Muller’s ratchet, proposing large-scale population genomics study for its validation, since this understanding could lead to therapeutic approaches for facilitating mutational meltdown of SARS-CoV-2, as experienced earlier for influenza A virus.

## Introduction

The emergence and rapid global spread of novel severe acute respiratory syndrome coronavirus 2 (SARS-CoV-2), the causative agent of the coronavirus disease 2019 (COVID-19), has led to an unprecedented worldwide public health crisis, crossing the species barrier and disseminating rapidly through the global population [1]. Like any other rapidly evolving emerging pathogens, effective preventive measures and treatment approaches call for an indepth understanding of the evolutionary dynamics of SARS-CoV-2 and its association with the epidemiological data [2, 3], especially with distinct attention to specific geographic regions. Nature and strength of selection pressures acting on the pathogen can vary across different ethnicities, countries, or even among specific administrative regions (such as states or provinces) within a country like India, China, USA, largely because of the implementation of social distancing via lockdown as a preventive measure against the geographic spread of the pathogen between and within countries.

The transmembrane spike glycoprotein protrudes from the viral surface and is responsible for viral attachment, fusion and entrance into the host cells, thereby establishing the infection [4, 5]. The two most notable genetic features of this protein are (a) the binding of its surface unit (S1) to the human cellular receptor angiotensin converting enzyme 2 (hACE2), and (b) the fusion of viral membrane with a host cell membrane via its transmembrane unit (S2). The S1 subunit contains the receptor-binding domains (RBD) supporting the stabilization of the membrane-anchored state of the S2 subunit along with its fusion machinery [6-11]. Some previous works have acknowledged that extensive and irreversible conformational changes stimulate the cleavage of spike protein, thereby activating it for membrane fusion [11-17]. Though it is not yet confirmed if the differences owing to conformational changes are aiding to the expansion of SAR-CoV2 varieties increasing or decreasing its infectivity or transmissibility, but the researchers confirmed the spike proteins to be the key pathogenic determinant that differentiates SAR-CoV2 from other SARS-related coronaviruses [17]. It is predicted that mutations in the spike protein increases or decreases protein glycosylation, thereby enhancing or reducing the viral uptake by host cells [18-22]. Although the RBD in the spike protein is known as the most important player to recognize the host receptor, it is highly likely that the region outside of this C-terminal domain in S1 subunit as well as the domains of S2 subunit also influence the host-receptor binding allosterically.

The present study analyses spike protein variants from Indian isolates to understand the evolutionary patterns and driving forces behind the emerging SARS-CoV-2 infection and its potential epidemiological footprints in India. We detected a strong correlation between the average stability of complexes formed by the circulating spike protein variants with the host receptor (S-R complex) and the disease severity of a given location, suggesting the S-R complex stability as a potential marker to assess the severity of the disease. Importantly, majority of the emerging variants showed decreased stability, indicating accumulation of deleterious mutations in the spike protein. This conforms to the declining fatality rates of the disease countrywide. Could the fixation of these deleterious mutations in the population lead to mutational meltdown, following Muller’s ratchet dynamics of evolution?

## Methods

### Analysis of sequence diversity and reconstruction of phylogeny

The average pairwise nucleotide diversity (π) and the rates of synonymous (dS) and nonsynonymous (dN) mutations for the spike protein-coding genes were calculated using MEGA version X [23]. TimeZone software [24] was used to reconstruct the maximumlikelihood based phylogeny to map the protein variants and identify convergent amino acid changes (i.e. repeated independent or phylogenetically unlinked mutations at the same amino acid positions). The spike gene sequence from Wuhan-Hu-1/2019 genome was used as reference to detect the orthologs in the sequenced Indian genomes based on a threshold value of 95% for both nucleotide sequence diversity and gene length coverage.

### Identification of spike protein variants unique to India

All 17529 spike protein sequences from worldwide isolates available till May 9, 2020 in the GISAID database (https://www.gisaid.org/) were downloaded. We implemented CD-HIT Suite [25, 26] to cluster all the spike protein sequences considering 100% amino acid sequence identity as ortholog clustering criteria, and detected a total of 3706 clusters. Of these clusters, we considered 3577 clusters matching the complete length of spike protein (1273 amino acids), using the sequence from Wuhan-Hu-1/2019 (GenBank accession number MN908947) as reference. These 3577 spike protein sequences were aligned using ClustalW program [27, 28]. The resulting alignment was compared with Indian mutational variants mapped in the previous step to distinguish the variants unique to Indian isolates.

### Analysis of state-wise diversity of spike protein variants

For each Indian state, we computed the number of spike protein variants and the frequency of each of those variants. The state-wise calculation of variant diversity was performed using Simpson’s index [29].

### Modeling of spike protein – hACE2 complex variants

Variants of the ancestral spike protein (Wuhan-Hu-1/2019) were built by homology modeling using Swiss modeler [30] with the aid of available templates (residue range: 27-1147) using Wuhan-Hu-1/2019 isolate as reference [17, 31]. X-ray crystal structure of the human ACE2 was used from the complex of receptor binding domain (RBD) of spike protein with hACE2 (PDB code: 6lzg) [32]. We docked the RBD (residue range: 331-524) of spike protein (residue range: 27-1147) mutants to the binding site of hACE2 using HADDOCK webserver [33, 34] by providing binding site information [31, 35]. HADDOCK score (often mentioned as docking score in the text), with some arbitrary unit, signifies a measure determined by weighted sum of intermolecular interactions, such as electrostatic and van der Waals interactions between protein and ligand, desolvation energy, restraint violation energy and the buried surface area upon binding. For each docked complex HADDOCK score was estimated and VMD [36] was used to visualize the structures.

## Results

### Two major ancestors circulating in India lead to a burst of spike protein variants

We identified a total of 630 isolates with complete gene sequences encoding the spike protein based on the submissions of Indian SARS CoV-2 genome sequences to GISAID till June 7, 2020 (S1 Table). The samples analyzed were isolated from 17 states and 2 union territories of India, collectively called as ‘states’ hereafter. We found a countrywide average pairwise nucleotide diversity (π) of 0.048±0.02%. The rates of synonymous (or silent) and nonsynonymous (or amino acid replacement) mutations were found to be 0.097±0.05% and 0.033±0.02% respectively. Phylogenetic analysis showed the Wuhan-Hu-1/2019 variant of the spike protein as the most ancestral one, as expected, while the D614G variant emerged from the Wuhan-Hu-1/2019 variant appeared to be another stable variant circulating in the Indian population. Since both these variants have established themselves in the worldwide population as two major ancestors of SARS CoV-2 spike protein variants, we here onwards will refer the Wuhan-Hu-1/2019 and D614G variants as ancestor 1 and ancestor 2 respectively.

Apart from giving rise to ancestor 2, the ancestor 1 led to a total of 20 variants (Fig 1 and S2 Table). Of these, the variant K77M evolved further to yield three more variants isolated from three different states (Bihar, Tamil Nadu and Telengana), suggesting the emergence of K77M as another stable variant. On the other hand, the ancestor 2 showed about twice more diversity by giving rise to 47 variants (Fig 1 and S2 Table). In this ancestor 2 clade, several variants (L5F, T22I, L54F, G261S, T572I, E583D, Q677H, A706S, H1083Q) indicated their stability in the population via mutating further, giving rise to additional variants. Besides, although the ancestral variants were predominant in the population circulating in India, we detected a total of 16 variants of spike proteins that were represented by multiple isolates (Fig 1), from 2 to as many as 9 isolates, indicating the possible fixation of some of these variants in the population irrespective of the stability of S-R complex. Of these, an array of mutations in 8 variants (at positions 5, 54, 78, 558, 574, 583, 677, 1243) showed their convergent nature, where those mutations at the same positions were phylogenetically unlinked, i.e. repeated independently (S2 Table). Interestingly, 53% of the total set of variants detected in Indian population was found unique, i.e. not found in 17529 worldwide genomic isolates analyzed from the GISAID database till May 09, 2020 (S2 Table). We believe that this considerable level of uniqueness could be an expected scenario in almost all geographical regions where a newly emerging, rapidly evolving viral pathogen tries to adapt to a new host, and many of these variants might be detrimental to the fitness of the organism. However, specific positive selection pressures could also play a role in this mutational pattern which needs to be studied separately in some greater depth.

**Fig. 1.**
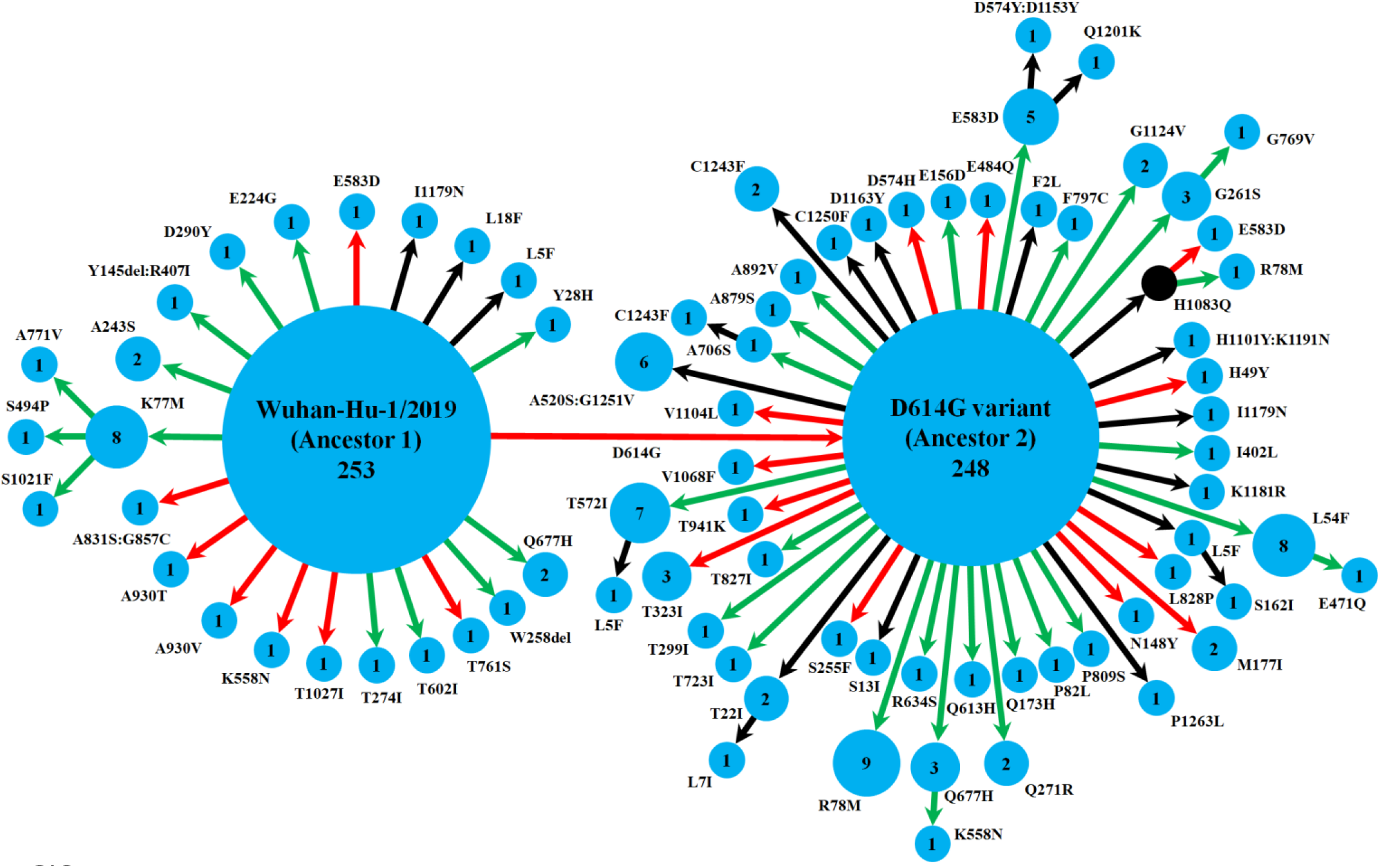
Schematic representation of the diversity of spike protein variants circulating in India, using maximum likelihood-based phylogeny reconstruction. Each node represents a specific spike protein variant, while the node-size and the number inside depict the frequency of that variant. The red or green color of each arrow indicates the higher or lower stability index respectively of the S-R complex for each variant than the major ancestral variant it emerged from (either Ancestor 1 or Ancestor 2). The black arrows lead to the variants for which the docking scores could not be determined either because of the presence of at least one variation outside the available template region for docking(18, 42), or due to non-existing isolate in the lone hypothetical node with H1083Q mutation denoted by black color. This black node signifies a variant with no available isolate in our dataset, while it gives rise to two derived variants, H1083Q:R78M and H1083Q:E583D, for which representative isolates were available.

### The average stability index of S-R complex correlates strongly with the fatality rates in a given location

We next looked into the distribution of these spike protein variants across Indian states (S3 Table). As we computed the diversity based on both richness and evenness of spike variants, some of the states like Maharashtra, Odisha, West Bengal and Gujarat demonstrated significantly higher (P<0.05) diversity of circulating variants than most of the remaining states. However, while we had 196, 97, 75 and 73 sequenced isolates from Gujarat, Telengana, Delhi and Maharashtra respectively, the remaining states were represented by even lower sample size (ranging from 1 to 48 isolates. Of these, Delhi variants showed lowest diversity, significantly different from both Maharashtra (P=0.0002) and Gujarat (P=0.0006), though not from Telengana (P=0.16). On the other hand, Maharashtra variants presented much higher diversity than Telengana (P=0.017) or Gujarat (P=0.21).

It is expected that these variations might be in response to strong selection pressures acting on the spike protein, especially its S1 subunit which, being a major immunogenic target for the host, plays the pivotal role to evade the host immune response and to offer a successful viral entry. Therefore, the mutational variations in spike proteins can essentially affect the stability of the S-R complex. To assess this, we modeled each of the spike protein variants, and then docked to the binding site of host receptor, hACE2, using HADDOCK webserver [33, 34] of data-driven docking algorithm by providing binding site information as the same was already established from the crystal structure [32, 35]. In the circulating variants, we detected a significant excess (P=0.035) of mutations in the S1 subunit (with 41 mutations) compared to the S2 subunit (with 22 mutations). The mutation positions in the secondary structures of the analyzed variants are detailed in the S1 Dataset in supporting information.

The docking score (HADDOCK score) of each variant on hACE2 is hereafter designated as the stability index of S-R complex. More negative is the docking (HADDOCK) score, higher is the stability of the S-R complex [37]. Under the assumption that better stability would lead to better invasion of the virus into the host, we hypothesize that such a stability index of a given spike protein variant could be linked to the severity of viral pathogenicity. To test this hypothesis, we measured the severity as the ‘fatality rate’ calculated simply as the ratio of the number of deceased people to the number of recovered in a given state (available from the Government of India website: www.mygov.in/corona-data/covid19-statewise-status/).

While we aimed to estimate an average stability index of a given state based on the stability indices of circulating variants in that location, we were handicapped with the available sample size and the information of collection diversity. Considering this issue, we restricted our study to the states having 50 or more sequenced samples for analysis. Therefore, we could assess the association of average stability index with fatality rates for three states and one union territory (Maharashtra, Gujarat, Telengana and Delhi) which qualified our sample size threshold. Importantly, these four regions harbored 70% of all samples analyzed across 19 states, while their variant diversity ranged from the highest to one of the lowest (S3 Table).

We detected a strong exponential correlation (R^2^=0.96) between the average stability index of circulating spike variants of the region with the fatality rate in that region (Fig 2). While Telengana and Delhi showed comparable average stability index values with ~7% fatality rates, Maharashtra and Gujarat had exponentially higher stability index values (i.e., more negative docking or HADDOCK scores) with 8% and 9% fatality rates respectively. It is highly plausible that the spike protein, as the primary controller of both the attachment to the host cell surface and the initiation of infection by fusing the viral and the host cell membranes, would be represented by variants with varying efficiency of the virus to enter human cells, and to get transmitted among people [17]. However, our conclusions based on the available initial data are premature due to low sample size per location and lack of direct evidence for the correlation between spike protein variant’s docking score and the pathogen’s contribution to host fatality, thereby warranting population-level robust association analysis and experimental validations.

**Fig. 2.**
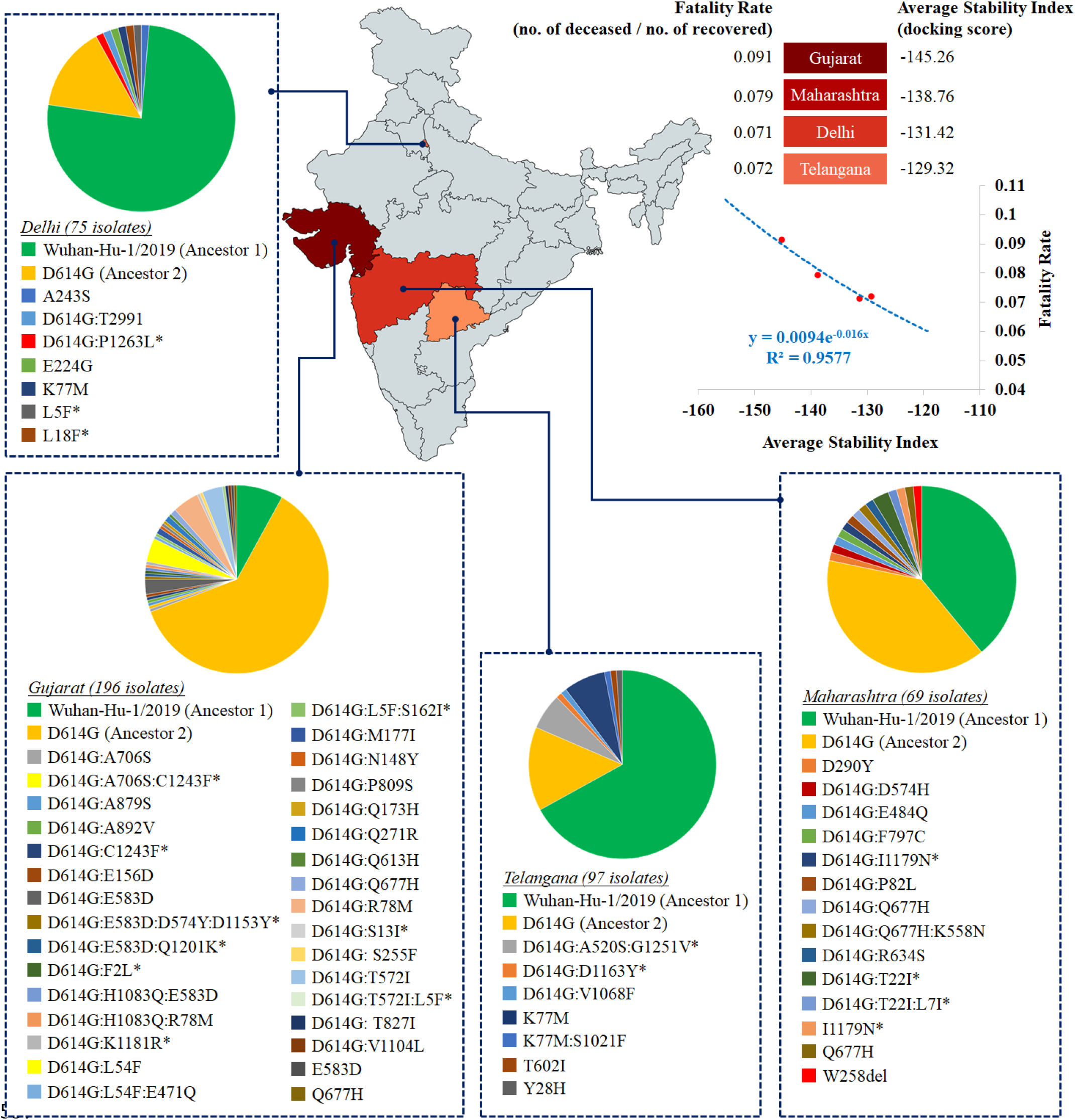
Heat map distribution across four Indian states with >50 sequenced isolates based on average stability index. The average stability index for a particular state denotes the averaged value of docking scores / HADDOCK scores of S-R complexes for all circulating variants. The values of average stability index and fatality rate in Indian states are plotted to fit an exponential function (R^2^=0.96).

### The emerging spike protein variants showing reduced stability of S-R complex are significantly abundant

Of 630 isolates analyzed, Ancestors 1 and 2 were represented by 253 and 248 isolates, suggesting their steady circulation across India. However, the remaining ones, i.e. more than 20% isolates represented relatively recently emerged variants out of two ancestral variants. Interestingly, a quick look at Fig 1 showed that, for majority of the variants derived from the Ancestor 1 and Ancestor 2, the stability was reduced (having less negative stability index values) relative to their respective ancestors. We therefore plotted the trend of those emerging variants with reference to their ancestral backgrounds (Fig 3). Significant majority (χ^2^ P=0.03) of the variants that emerged from the two ancestral variants showed reduced stability (having less negative docking scores) from their respective ancestors. This picture got even more prominent (χ^2^ P=0.009) when we looked into exclusively the variants which were detected multiple times in the dataset, i.e. represented by more than one isolate (sometimes collected from different states), thereby suggesting possible fixation of those variants in the population (S2 Table). Interestingly, 13 of these variants with multiple occurrences accumulated mutations exclusively in S1 subunit, while only 3 variants showed all mutations in S2 subunit, which might be suggesting an increased selection pressure in the S1 subunit because of its key role in the viral entry and the presence of the receptor binding domain.

**Fig. 3.**
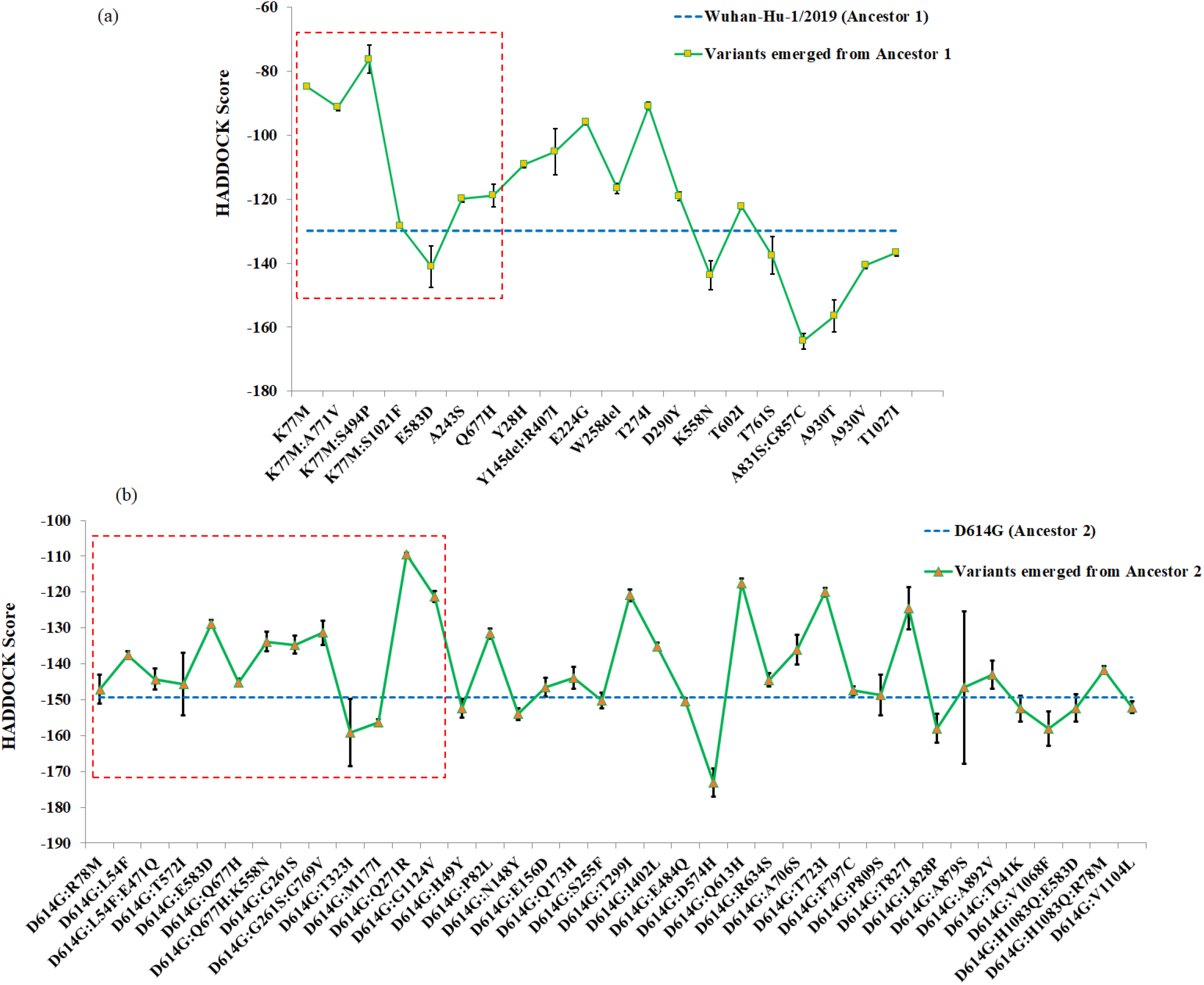
Stability index (i.e., docking score or HADDOCK score) plot of S-R complexes for spike protein variants emerging from (a) Ancestor 1 (Wuhan-Hu-1/2019 variant) and (b) Ancestor 2 (D614G variant). The blue dotted line is used as a reference to denote the stability index value for respective ancestral variants. More negative is the value, higher is the stability level. The red dotted rectangular block includes the variants that are represented by multiple isolates in our dataset (*SI appendix*, Table S2).

At this point, we can propose that the relatively recent variants emerging from the ancestors in India are losing their ability on an average to form a stable complex with the human receptor as compared with their ancestors, which could possibly result in lower countrywide fatality rate, if we combine our earlier observation of the direct correlation between average stability index and fatality rate. Conforming to this proposal, we found that, after an initial steady increase of fatality rate for the first four weeks, the fatality rate reached a peak at 38.2% on April 11, 2020, followed by a continuous sharp decline at 5.3% on June 28, 2020 until when the data were available (Fig 4), fitting a power law function (R^2^=0.94) (data source: http://covidindiaupdates.in/). This was despite the fact that the rate of mutations was found to be increasing during the three-month period from March till May 2020 during which our analyzed samples were collected. Interestingly, the increase in nonsynonymous (i.e. amino acid replacement) mutation rate (from 0.027% in March to 0.033% in May 2020) was detected to be about 14% higher than the increase in synonymous mutation rate (from 0.091% in March to 0.097% in May 2020), suggesting a stronger selection pressure for amino acid changes in the spike protein.

**Fig. 4.**
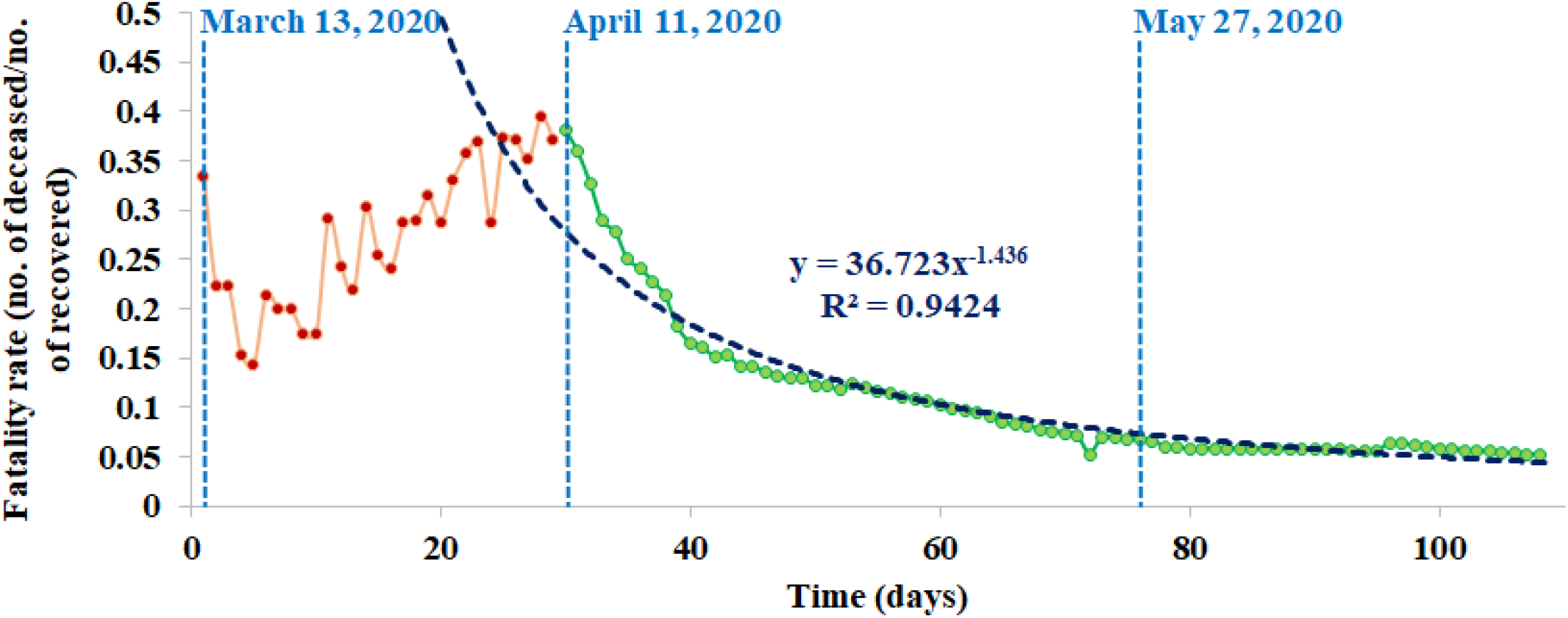
Countrywide fatality rates (the number of deaths / the number of recovered cases) over time in India. The date of first available data (March 13, 2020), the date after which the decline of fatality rate started (April 11, 2020), and the date until which the analyzed samples were collected (May 27, 2020) are denoted by blue dotted lines. The declining fatality rate curve is best fitted by a power law function (R^2^=0.94).

### Could Muller’s ratchet be a player in shaping SARS-CoV-2 evolutionary dynamics in India?

In evolutionary genetics, Muller’s ratchet signifies accumulation of deleterious mutation in a population leading to ‘mutational meltdown’, leading to a gradual extinction of that population [38]. In general, genetic mutations that provide adaptive advantages are fixed in the population by natural selection whereas deleterious ones are wiped off from the population. However, an accelerated mutation rate puts huge mutational pressure, and natural selection is unable to wash out these deleterious mutations, retaining the newly formed variants within the population and thereby leading to their fixation. This irretrievable evolutionary mechanism is coined as Muller’s ratchet by evolutionary biologists [39]. When more and more deleterious mutations are accumulated and become permanent in the population, this results in ‘mutational meltdown’ or ultimate loss of the population [40, 41].

It is known that the apparent tendency to directly correlate the high mutation rate of virus with its infectivity and transmissibility is without merit [42]. On the other side, it has been suggested that Muller’s ratchet, via mutational meltdown, could be a key player in leading the SARS-CoV-2 population to gradual extinction due to accumulation and fixation of deleterious mutations in future [40, 43]. On one hand, we observed that the stability of S-R complex is directly linked to the fatality rates, while the continuous emergence of variants from the ancestral ones were found to be less stable compared to their ancestors. As we combine these findings with the countrywide data of a sharp decline in the fatality rate over time, we propose the possibility of mutational meltdown in action for SARS-CoV-2 in India, indicating Muller’s ratchet as a plausible game-changer for COVID-19 scenario here in near future.

We understand that, only in three months, we cannot expect a radical increase in the mutation rate to give rise to any significant accumulation of deleterious mutations that could have offered a prominent picture of mutational meltdown. However, our results altogether point toward the trend, thereby suggesting the potential of future studies in this otherwise overlooked domain of microbial dynamics, which could in turn lead to a possibility of a successful therapeutic approach [40, 43].

## Discussion

Our work has combined genetic and epidemiological data of SARS-CoV2 in India to decipher a direct correlation between the average stability of S-R complexes for the circulating spike protein variants and the fatality rate of a geographic region. The docking score of S-R complex, designated here as the stability index, is estimated by protein-protein docking based on intermolecular interactions, such as electrostatic and van der Waals interactions, desolvation and restraint violation energies along with buried surface area upon binding via detection of the correct binding pose. This score, in essence, quantifies the stability of the docked complex by optimizing global minimum energy conformation of the complex [44, 45]. On the other side, while it is probable that a better stability of S-R complex could lead to an increased viral load and maybe an increased infectivity, previous works showed that the spike protein – hACE2 complex is crucial for viral pathogenesis by causing acute lung damage [46, 47], which suggests a direct link between S-R complex stability and fatality rate. However, the robustness of the potential of S-R complex stability index for the spike protein variants as a tracker of fatality rate or disease severity needs to be studied in greater depths with more structured region-specific patient data (that are primarily available to selected government/non-government agencies) in connection with larger population-level sequence datasets for the given locations.

As expected for any fast-moving pathogen outbreaks [42], we find an increasing rate of mutations, while the extent of increase is much higher for the nonsynonymous (amino acid replacement) changes. Interestingly, the S-R complex of a significant majority of variants tend to lose stability relative to their two most stable ancestral variants. Alongside, the countrywide data show a continuous sharp decline in the fatality rate after an initial surge. Therefore, keeping in mind the observed correlation between the S-R complex stability and fatality rate across Indian states, here we pose an open question for future research: Does the phylodynamics of SARS-CoV-2 in India indicate any nascent action of Muller’s ratchet where the otherwise deleterious mutations tend to get fixed in the population as natural selection remains unable to purge them due to excessive mutational pressures? If so, there lies an immense potential of using therapeutics that could facilitate such a process of mutational meltdown, as was demonstrated earlier for influenza A virus [48, 49]. Therefore, future large-scale population genomics analysis supported by epidemiological information is of high importance to explore this question for India as well as for other countries across the globe to develop efficient analytical methods, thereby guiding better surveillance programs, prevention and treatment management of COVID-19.

## Supporting information Captions

**S1 Table:** List of the sequenced isolates analyzed from GISAID database (www.gisaid.org) available till June 7, 2020. These isolates included complete gene sequence for the spike protein. The mutations of spike protein variant for each isolate was mentioned along with the docking score of S-R complex for the variant. The docking scores of the variants with mutations between amino acid positions 1-26and 1148-1273 were not determined because of the unavailability of the modeling template (1, 2) and therefore mentioned as NA (not available).

**S2 Table:** Countrywide and state-wise frequency distribution matrix of circulating variants along with the docking score of S-R complex for each variant. The docking scores of the variants with mutations between amino acid positions 1-26 and 1148-1273 were not determined because of the unavailability of the modeling template (1, 2) and therefore mentioned as NA (not available). The variant mutation positions marked by asterisks denote convergent nature of these mutations, suggesting their independent accumulation multiple times along phylogenetically unlinked allelic backgrounds. The variants that were detected to be uniquely present in Indian population as compared to a total of 17529 worldwide genomic isolates available from the GISAID database till May 9, 2020.

**S3 Table:** State-wise distribution of the type and frequency of spike protein variants, along with the calculated diversity based on Simpson’s index (3).

**Dataset S1:** Positional information of the spike protein mutations in the docked S-R complex.

