## Supplemental Tables and Text for "Spike protein mutational landscape in India: Could Muller’s ratchet be a future game-changer for COVID-19?"

**Supporting Information**

**S1 Table.**

List of the sequenced isolates analyzed from GISAID database (www.gisaid.org) available till June 7, 2020. These isolates included complete gene sequence for the spike protein. The mutations of spike protein variant for each isolate was mentioned along with the docking score of S-R complex for the variant. The docking scores of the variants with mutations between amino acid positions 1-26 and 1148-1273 were not determined because of the unavailability of the modeling template [1, 2] and therefore mentioned as NA (not available).

| Accession ID | Virus name | Location | Collection date | Mutation Type | Docking Score | Standard Error |
|---|---|---|---|---|---|---|
| EPI_ISL_413522 | hCoV-19/India/1-27/2020 | Asia / India / Kerala | 2020-01-27 | Y145del:R407I | -105.2 | 7.245 |
| EPI_ISL_413523 | hCoV-19/India/1-31/2020 | Asia / India / Kerala | 2020-01-31 | A930V | -140.7 | 1.039 |
| EPI_ISL_426414 | hCoV-19/India/GBRC1/2020 | Asia / India / Gujarat | 2020-04-05 | D614G:Q271R | -109.5 | 0.477 |
| EPI_ISL_426415 | hCoV-19/India/GBRC1s/2020 | Asia / India / Gujarat | 2020-04-05 | D614G:Q271R | -109.5 | 0.477 |
| EPI_ISL_428479 | hCoV-19/India/nimh-0113/2020 | Asia / India / Karnataka | 2020-04-06 | D614G | -149.3 | 1.037 |
| EPI_ISL_428481 | hCoV-19/India/nimh-0130/2020 | Asia / India / Karnataka | 2020-04-06 | D614G | -149.3 | 1.037 |
| EPI_ISL_428482 | hCoV-19/India/nimh-0182/2020 | Asia / India / Karnataka | 2020-04-08 | D614G:C1250F | NA | NA |

| | | | | | | |
|---|---|---|---|---|---|---|
| EPI_ISL_428483 | hCoV-19/India/nimh-0318/2020 | Asia / India / Karnataka | 2020-04-10 | Ancestral | -129.8 | 1.124 |
| EPI_ISL_428484 | hCoV-19/India/nimh-0351/2020 | Asia / India / Karnataka | 2020-04-10 | Ancestral | -129.8 | 1.124 |
| EPI_ISL_428486 | hCoV-19/India/nimh-0996/2020 | Asia / India / Karnataka | 2020-04-14 | Ancestral | -129.8 | 1.124 |
| EPI_ISL_428487 | hCoV-19/India/nimh-1071/2020 | Asia / India / Karnataka | 2020-04-14 | Ancestral | -129.8 | 1.124 |
| EPI_ISL_430464 | hCoV-19/India/S3/2020 | Asia / India / West Bengal / Kolkata | 2020-03-21 | D614G:G1124V | -121.3 | 1.565 |
| EPI_ISL_430465 | hCoV-19/India/S5/2020 | Asia / India / West Bengal / Darjeeling | 2020-03-28 | D614G | -149.3 | 1.037 |
| EPI_ISL_430466 | hCoV-19/India/S6/2020 | Asia / India / West Bengal / Tehatta | 2020-03-26 | D614G:T723I | -120 | 1.201 |
| EPI_ISL_430467 | hCoV-19/India/S11/2020 | Asia / India / West Bengal / East Medinipur | 2020-04-03 | D614G | -149.3 | 1.037 |
| EPI_ISL_430468 | hCoV-19/India/S2/2020 | Asia / India / West_Bengal / Kolkata | 2020-03-21 | D614G:G1124V | -121.3 | 1.565 |
| EPI_ISL_431101 | hCoV-19/India/GMC-RK100/2020 | Asia / India / Telangana / Hyderabad | 2020-03-01 | Y28H | -109.2 | 1.092 |
| EPI_ISL_431102 | hCoV-19/India/GMC-KN318/2020 | Asia / India / Telangana / Hyderabad | 2020-03-11 | D614G | -149.3 | 1.037 |

| | | | | | | |
|---|---|---|---|---|---|---|
| EPI_ISL_431103 | hCoV-19/India/GMC-KN443/2020 | Asia / India / Telangana / Hyderabad | 2020-03-16 | Ancestral | -129.8 | 1.124 |
| EPI_ISL_431117 | hCoV-19/India/GMC-TC469/2020 | Asia / India / Telangana / Hyderabad | 2020-03-20 | D614G | -149.3 | 1.037 |
| EPI_ISL_435049 | hCoV-19/India/GBRC2/2020 | Asia / India / Gujarat / Ahmedabad | 2020-04-13 | D614G | -149.3 | 1.037 |
| EPI_ISL_435050 | hCoV-19/India/GBRC3/2020 | Asia / India / Gujarat / Ahmedabad | 2020-04-13 | D614G | -149.3 | 1.037 |
| EPI_ISL_435051 | hCoV-19/India/GBRC4/2020 | Asia / India / Gujarat / Ahmedabad | 2020-04-13 | D614G | -149.3 | 1.037 |
| EPI_ISL_435052 | hCoV-19/India/GBRC5/2020 | Asia / India / Gujarat / Ahmedabad | 2020-04-13 | D614G | -149.3 | 1.037 |
| EPI_ISL_435053 | hCoV-19/India/GBRC6/2020 | Asia / India / Gujarat / Ahmedabad | 2020-04-07 | D614G | -149.3 | 1.037 |
| EPI_ISL_435054 | hCoV-19/India/GBRC7/2020 | Asia / India / Gujarat / Ahmedabad | 2020-04-14 | D614G | -149.3 | 1.037 |
| EPI_ISL_435055 | hCoV-19/India/GBRC8/2020 | Asia / India / Gujarat / Gandhinagar | 2020-04-22 | D614G | -149.3 | 1.037 |
| EPI_ISL_435056 | hCoV-19/India/GBRC9/2020 | Asia / India / Gujarat / Mansa | 2020-04-21 | D614G | -149.3 | 1.037 |

| | | | | | | |
|---|---|---|---|---|---|---|
| EPI_ISL_435060 | hCoV-19/India/NCDC-01257/2020 | Asia / India / Uttar Pradesh / Noida | 2020-03-12 | D614G | -149.3 | 1.037 |
| EPI_ISL_435061 | hCoV-19/India/NCDC-01537/2020 | Asia / India / Delhi | 2020-03-16 | D614G:P1263L | NA | NA |
| EPI_ISL_435062 | hCoV-19/India/NCDC-01538/2020 | Asia / India / Punjab | 2020-03-16 | D614G | -149.3 | 1.037 |
| EPI_ISL_435063 | hCoV-19/India/NCDC-01326/2020 | Asia / India / Delhi | 2020-03-13 | D614G | -149.3 | 1.037 |
| EPI_ISL_435064 | hCoV-19/India/NCDC-01705/2020 | Asia / India / Delhi | 2020-03-18 | D614G | -149.3 | 1.037 |
| EPI_ISL_435065 | hCoV-19/India/NCDC-01475/2020 | Asia / India / Delhi | 2020-03-15 | D614G | -149.3 | 1.037 |
| EPI_ISL_435066 | hCoV-19/India/NCDC-01711/2020 | Asia / India / Delhi | 2020-03-18 | D614G | -149.3 | 1.037 |
| EPI_ISL_435067 | hCoV-19/India/NCDC-01712/2020 | Asia / India / Delhi | 2020-03-18 | D614G:T299I | -120.9 | 1.636 |
| EPI_ISL_435068 | hCoV-19/India/NCDC-01757/2020 | Asia / India / Delhi | 2020-03-18 | D614G | -149.3 | 1.037 |
| EPI_ISL_435069 | hCoV-19/India/NCDC-01744/2020 | Asia / India / Delhi | 2020-03-18 | D614G | -149.3 | 1.037 |
| EPI_ISL_435070 | hCoV-19/India/NCDC-01710/2020 | Asia / India / Delhi | 2020-03-18 | D614G | -149.3 | 1.037 |
| EPI_ISL_435071 | hCoV-19/India/NCDC-02105/2020 | Asia / India / Delhi | 2020-03-25 | D614G | -149.3 | 1.037 |
| EPI_ISL_435072 | hCoV-19/India/NCDC-02155/2020 | Asia / India / Delhi | 2020-03-26 | D614G | -149.3 | 1.037 |

| | | | | | | |
|---|---|---|---|---|---|---|
| EPI_ISL_435074 | hCoV-19/India/NCDC-02250/2020 | Asia / India / West Bengal | 2020-03-28 | Ancestral | -129.8 | 1.124 |
| EPI_ISL_435075 | hCoV-19/India/NCDC-02331/2020 | Asia / India / Tamil Nadu | 2020-03-29 | K77M:A771V | -91.3 | 1.125 |
| EPI_ISL_435078 | hCoV-19/India/NCDC-02311/2020 | Asia / India / Tamil Nadu | 2020-03-29 | Ancestral | -129.8 | 1.124 |
| EPI_ISL_435079 | hCoV-19/India/NCDC-02318/2020 | Asia / India / Tamil Nadu | 2020-03-29 | Ancestral | -129.8 | 1.124 |
| EPI_ISL_435080 | hCoV-19/India/NCDC-02334/2020 | Asia / India / Tamil Nadu | 2020-03-29 | Ancestral | -129.8 | 1.124 |
| EPI_ISL_435081 | hCoV-19/India/NCDC-02248/2020 | Asia / India / West Bengal | 2020-03-28 | Ancestral | -129.8 | 1.124 |
| EPI_ISL_435082 | hCoV-19/India/NCDC-02322/2020 | Asia / India / Uttar Pradesh | 2020-03-29 | Ancestral | -129.8 | 1.124 |
| EPI_ISL_435083 | hCoV-19/India/NCDC-02242/2020 | Asia / India / Tamil Nadu | 2020-03-28 | Ancestral | -129.8 | 1.124 |
| EPI_ISL_435084 | hCoV-19/India/NCDC-02309/2020 | Asia / India / Tamil Nadu | 2020-03-29 | Ancestral | -129.8 | 1.124 |
| EPI_ISL_435085 | hCoV-19/India/NCDC-02310/2020 | Asia / India / Mumbai | 2020-03-29 | Ancestral | -129.8 | 1.124 |
| EPI_ISL_435086 | hCoV-19/India/NCDC-02315/2020 | Asia / India / Mumbai | 2020-03-29 | Ancestral | -129.8 | 1.124 |
| EPI_ISL_435087 | hCoV-19/India/NCDC-02333/2020 | Asia / India / Tamil Nadu | 2020-03-29 | Ancestral | -129.8 | 1.124 |
| EPI_ISL_435088 | hCoV-19/India/NCDC-02336/2020 | Asia / India / Odisha | 2020-03-29 | Ancestral | -129.8 | 1.124 |
| EPI_ISL_435090 | hCoV-19/India/NCDC-02326/2020 | Asia / India / Jammu | 2020-03-29 | Ancestral | -129.8 | 1.124 |

| | | | | | | |
|---|---|---|---|---|---|---|
| EPI_ISL_435091 | hCoV-19/India/NCDC-02245/2020 | Asia / India / Tamil Nadu | 2020-03-28 | Ancestral | -129.8 | 1.124 |
| EPI_ISL_435092 | hCoV-19/India/NCDC-02244/2020 | Asia / India / Tamil Nadu | 2020-03-28 | Ancestral | -129.8 | 1.124 |
| EPI_ISL_435093 | hCoV-19/India/NCDC-02312/2020 | Asia / India / Tamil Nadu | 2020-03-29 | Ancestral | -129.8 | 1.124 |
| EPI_ISL_435094 | hCoV-19/India/NCDC-02327/2020 | Asia / India / Tamil Nadu | 2020-03-29 | Ancestral | -129.8 | 1.124 |
| EPI_ISL_435095 | hCoV-19/India/NCDC-02328/2020 | Asia / India / Tamil Nadu | 2020-03-29 | Ancestral | -129.8 | 1.124 |
| EPI_ISL_435096 | hCoV-19/India/NCDC-02332/2020 | Asia / India / Tamil Nadu | 2020-03-29 | Ancestral | -129.8 | 1.124 |
| EPI_ISL_435097 | hCoV-19/India/NCDC-02252/2020 | Asia / India / West Bengal | 2020-03-28 | Ancestral | -129.8 | 1.124 |
| EPI_ISL_435099 | hCoV-19/India/NCDC-02320/2020 | Asia / India / Uttar Pradesh | 2020-03-29 | Ancestral | -129.8 | 1.124 |
| EPI_ISL_435100 | hCoV-19/India/NCDC-02321/2020 | Asia / India / Uttar Pradesh | 2020-03-29 | Ancestral | -129.8 | 1.124 |
| EPI_ISL_435101 | hCoV-19/India/NCDC-01441/2020 | Asia / India / Ladakh | 2020-03-15 | Ancestral | -129.8 | 1.124 |
| EPI_ISL_435102 | hCoV-19/India/NCDC-01444/2020 | Asia / India / Ladakh | 2020-03-15 | Ancestral | -129.8 | 1.124 |
| EPI_ISL_435103 | hCoV-19/India/NCDC-01604/2020 | Asia / India / Ladakh | 2020-03-17 | Ancestral | -129.8 | 1.124 |
| EPI_ISL_435104 | hCoV-19/India/NCDC-01614/2020 | Asia / India / Ladakh | 2020-03-17 | Ancestral | -129.8 | 1.124 |
| EPI_ISL_435105 | hCoV-19/India/NCDC-01616/2020 | Asia / India / Ladakh | 2020-03-17 | Ancestral | -129.8 | 1.124 |

| | | | | | | |
|---|---|---|---|---|---|---|
| EPI_ISL_435106 | hCoV-19/India/NCDC-01760/2020 | Asia / India / Ladakh | 2020-03-18 | Ancestral | -129.8 | 1.124 |
| EPI_ISL_435107 | hCoV-19/India/NCDC-02370/2020 | Asia / India / Kargil | 2020-03-29 | Ancestral | -129.8 | 1.124 |
| EPI_ISL_435108 | hCoV-19/India/NCDC-01457/2020 | Asia / India / Delhi | 2020-03-15 | L5F | NA | NA |
| EPI_ISL_435109 | hCoV-19/India/NCDC-01638/2020 | Asia / India / Delhi | 2020-03-17 | Ancestral | -129.8 | 1.124 |
| EPI_ISL_435110 | hCoV-19/India/NCDC-01501/2020 | Asia / India / Delhi | 2020-03-16 | L18F | NA | NA |
| EPI_ISL_435111 | hCoV-19/India/NCDC-02415/2020 | Asia / India / Delhi | 2020-03-30 | Ancestral | -129.8 | 1.124 |
| EPI_ISL_435112 | hCoV-19/India/NCDC-02251/2020 | Asia / India / Bihar | 2020-03-28 | Ancestral | -129.8 | 1.124 |
| EPI_ISL_436137 | hCoV-19/India/nimh-2873/2020 | Asia / India / Karnataka | 2020-04-21 | Ancestral | -129.8 | 1.124 |
| EPI_ISL_436138 | hCoV-19/India/nimh-3952/2020 | Asia / India / Karnataka | 2020-04-23 | D614G | -149.3 | 1.037 |
| EPI_ISL_436139 | hCoV-19/India/nimh-3970/2020 | Asia / India / Karnataka | 2020-04-23 | D614G | -149.3 | 1.037 |
| EPI_ISL_436140 | hCoV-19/India/nimh-4376/2020 | Asia / India / Karnataka | 2020-04-24 | D614G | -149.3 | 1.037 |
| EPI_ISL_436141 | hCoV-19/India/nimh-4378/2020 | Asia / India / Karnataka | 2020-04-24 | D614G | -149.3 | 1.037 |
| EPI_ISL_436156 | hCoV-19/India/nimh-1596/2020 | Asia / India / Karnataka | 2020-04-17 | Ancestral | -129.8 | 1.124 |
| EPI_ISL_436157 | hCoV-19/India/nimh-1598/2020 | Asia / India / Karnataka | 2020-04-17 | Ancestral | -129.8 | 1.124 |

| | | | | | | |
|---|---|---|---|---|---|---|
| EPI_ISL_436413 | hCoV-19/India/NCDC-2443/2020 | Asia / India / Uttar Pradesh | 2020-03-30 | D614G | -149.3 | 1.037 |
| EPI_ISL_436414 | hCoV-19/India/NCDC-2502/2020 | Asia / India / West Bengal | 2020-03-31 | Ancestral | -129.8 | 1.124 |
| EPI_ISL_436415 | hCoV-19/India/NCDC-2507/2020 | Asia / India / Delhi | 2020-03-31 | Ancestral | -129.8 | 1.124 |
| EPI_ISL_436416 | hCoV-19/India/NCDC-2514/2020 | Asia / India / Tamil Nadu | 2020-03-31 | T274I | -90.9 | 1.283 |
| EPI_ISL_436417 | hCoV-19/India/NCDC-2515/2020 | Asia / India / Bihar | 2020-03-31 | Ancestral | -129.8 | 1.124 |
| EPI_ISL_436418 | hCoV-19/India/NCDC-2517/2020 | Asia / India / Tamil Nadu | 2020-03-31 | Ancestral | -129.8 | 1.124 |
| EPI_ISL_436419 | hCoV-19/India/NCDC-2519/2020 | Asia / India / Bihar | 2020-03-31 | K77M:S494P | -76.3 | 4.382 |
| EPI_ISL_436420 | hCoV-19/India/NCDC-2522/2020 | Asia / India / Rajasthan | 2020-03-31 | Ancestral | -129.8 | 1.124 |
| EPI_ISL_436421 | hCoV-19/India/NCDC-2525/2020 | Asia / India / Assam | 2020-03-31 | Ancestral | -129.8 | 1.124 |
| EPI_ISL_436422 | hCoV-19/India/NCDC-2535/2020 | Asia / India / Assam | 2020-03-31 | Ancestral | -129.8 | 1.124 |
| EPI_ISL_436424 | hCoV-19/India/NCDC-3163/2020 | Asia / India / Delhi | 2020-04-05 | Ancestral | -129.8 | 1.124 |
| EPI_ISL_436425 | hCoV-19/India/NCDC-3170/2020 | Asia / India / Delhi | 2020-04-05 | Ancestral | -129.8 | 1.124 |
| EPI_ISL_436426 | hCoV-19/India/NCDC-3175/2020 | Asia / India / Delhi | 2020-04-05 | A243S | -119.9 | 0.950 |
| EPI_ISL_436427 | hCoV-19/India/NCDC-3176/2020 | Asia / India / Delhi | 2020-04-05 | Ancestral | -129.8 | 1.124 |

| | | | | | | |
|---|---|---|---|---|---|---|
| EPI_ISL_436428 | hCoV-19/India/NCDC-3181/2020 | Asia / India / Delhi | 2020-04-05 | Ancestral | -129.8 | 1.124 |
| EPI_ISL_436429 | hCoV-19/India/NCDC-3190/2020 | Asia / India / Delhi | 2020-04-05 | Ancestral | -129.8 | 1.124 |
| EPI_ISL_436430 | hCoV-19/India/NCDC-3195/2020 | Asia / India / Delhi | 2020-04-05 | Ancestral | -129.8 | 1.124 |
| EPI_ISL_436431 | hCoV-19/India/NCDC-3230/2020 | Asia / India / Delhi | 2020-04-05 | Ancestral | -129.8 | 1.124 |
| EPI_ISL_436432 | hCoV-19/India/NCDC-3264/2020 | Asia / India / Delhi | 2020-04-05 | Ancestral | -129.8 | 1.124 |
| EPI_ISL_436433 | hCoV-19/India/NCDC-3271/2020 | Asia / India / Delhi | 2020-04-05 | Ancestral | -129.8 | 1.124 |
| EPI_ISL_436434 | hCoV-19/India/NCDC-3279/2020 | Asia / India / Delhi | 2020-04-05 | Ancestral | -129.8 | 1.124 |
| EPI_ISL_436435 | hCoV-19/India/NCDC-3394/2020 | Asia / India / Delhi | 2020-04-09 | Ancestral | -129.8 | 1.124 |
| EPI_ISL_436436 | hCoV-19/India/NCDC-3400/2020 | Asia / India / Delhi | 2020-04-06 | Ancestral | -129.8 | 1.124 |
| EPI_ISL_436437 | hCoV-19/India/NCDC-3831/2020 | Asia / India / Delhi | 2020-04-08 | E224G | -95.9 | 0.857 |
| EPI_ISL_436438 | hCoV-19/India/NCDC-3835/2020 | Asia / India / Maharashtra | 2020-04-07 | Ancestral | -129.8 | 1.124 |
| EPI_ISL_436439 | hCoV-19/India/NCDC-3934/2020 | Asia / India / Bihar | 2020-04-09 | Ancestral | -129.8 | 1.124 |
| EPI_ISL_436440 | hCoV-19/India/NCDC-3936/2020 | Asia / India / Andhra Pradesh | 2020-04-09 | Ancestral | -129.8 | 1.124 |

| | | | | | | |
|---|---|---|---|---|---|---|
| EPI_ISL_436441 | hCoV-19/India/NCDC-3941/2020 | Asia / India / Bihar | 2020-04-09 | Ancestral | -129.8 | 1.124 |
| EPI_ISL_436442 | hCoV-19/India/NCDC-3950/2020 | Asia / India / Maharashtra | 2020-04-09 | D290Y | -119.1 | 1.389 |
| EPI_ISL_436443 | hCoV-19/India/NCDC-3961/2020 | Asia / India / Madhya Pradesh | 2020-04-09 | Ancestral | -129.8 | 1.124 |
| EPI_ISL_436444 | hCoV-19/India/NCDC-3965/2020 | Asia / India / Maharashtra | 2020-04-09 | Ancestral | -129.8 | 1.124 |
| EPI_ISL_436445 | hCoV-19/India/NCDC-3982/2020 | Asia / India / Delhi | 2020-04-09 | Ancestral | -129.8 | 1.124 |
| EPI_ISL_436447 | hCoV-19/India/NCDC-4055/2020 | Asia / India / Karnataka | 2020-04-10 | Ancestral | -129.8 | 1.124 |
| EPI_ISL_436448 | hCoV-19/India/NCDC-4138/2020 | Asia / India / Delhi | 2020-04-12 | Ancestral | -129.8 | 1.124 |
| EPI_ISL_436449 | hCoV-19/India/NCDC-4155/2020 | Asia / India / Bihar | 2020-04-12 | A243S | -119.9 | 0.950 |
| EPI_ISL_436450 | hCoV-19/India/NCDC-4189/2020 | Asia / India / Delhi | 2020-04-13 | K77M | -84.8 | 0.194 |
| EPI_ISL_436451 | hCoV-19/India/NCDC-4205/2020 | Asia / India / Delhi | 2020-04-13 | Ancestral | -129.8 | 1.124 |
| EPI_ISL_436452 | hCoV-19/India/NCDC-4208/2020 | Asia / India / Delhi | 2020-04-14 | Ancestral | -129.8 | 1.124 |
| EPI_ISL_436454 | hCoV-19/India/NCDC-4475/2020 | Asia / India / Delhi | 2020-04-13 | D614G | -149.3 | 1.037 |
| EPI_ISL_436455 | hCoV-19/India/NCDC-4476/2020 | Asia / India / Delhi | 2020-04-13 | D614G | -149.3 | 1.037 |

| | | | | | | |
|---|---|---|---|---|---|---|
| EPI_ISL_436456 | hCoV-19/India/NCDC-4846/2020 | Asia / India / Madhya Pradesh | 2020-04-20 | D614G | -149.3 | 1.037 |
| EPI_ISL_436457 | hCoV-19/India/NCDC-4859/2020 | Asia / India / Madhya Pradesh | 2020-04-20 | D614G | -149.3 | 1.037 |
| EPI_ISL_436459 | hCoV-19/India/NCDC-4874/2020 | Asia / India / Madhya Pradesh | 2020-04-20 | D614G | -149.3 | 1.037 |
| EPI_ISL_436460 | hCoV-19/India/NCDC-4876/2020 | Asia / India / Madhya Pradesh | 2020-04-20 | D614G | -149.3 | 1.037 |
| EPI_ISL_436461 | hCoV-19/India/NCDC-4877/2020 | Asia / India / Madhya Pradesh | 2020-04-20 | D614G | -149.3 | 1.037 |
| EPI_ISL_436462 | hCoV-19/India/NCDC-4878/2020 | Asia / India / Madhya Pradesh | 2020-04-16 | D614G | -149.3 | 1.037 |
| EPI_ISL_437440 | hCoV-19/India/GBRC12/2020 | Asia / India / Gujarat / Surat | 2020-04-26 | D614G:K1181R | NA | NA |
| EPI_ISL_437441 | hCoV-19/India/GBRC13/2020 | Asia / India / Gujarat / Surat | 2020-04-26 | Ancestral | -129.8 | 1.124 |
| EPI_ISL_437442 | hCoV-19/India/GBRC14/2020 | Asia / India / Gujarat / Surat | 2020-04-27 | Ancestral | -129.8 | 1.124 |
| EPI_ISL_437444 | hCoV-19/India/GBRC16/2020 | Asia / India / Gujarat / Surat | 2020-04-26 | Ancestral | -129.8 | 1.124 |
| EPI_ISL_437445 | hCoV-19/India/GBRC17a/2020 | Asia / India / Gujarat / Ahmedabad | 2020-04-26 | D614G:A706S:C1243F | NA | NA |

| | | | | | | |
|---|---|---|---|---|---|---|
| EPI_ISL_437446 | hCoV-19/India/GBRC17b/2020 | Asia / India / Gujarat / Ahmedabad | 2020-04-26 | D614G:C1243F | NA | NA |
| EPI_ISL_437447 | hCoV-19/India/GBRC18/2020 | Asia / India / Gujarat / Ahmedabad | 2020-04-26 | D614G | -149.3 | 1.037 |
| EPI_ISL_437448 | hCoV-19/India/GBRC19/2020 | Asia / India / Gujarat / Ahmedabad | 2020-04-26 | D614G | -149.3 | 1.037 |
| EPI_ISL_437449 | hCoV-19/India/GBRC20/2020 | Asia / India / Gujarat / Ahmedabad | 2020-04-26 | D614G | -149.3 | 1.037 |
| EPI_ISL_437450 | hCoV-19/India/GBRC21/2020 | Asia / India / Gujarat / Ahmedabad | 2020-04-26 | D614G:Q677H | -145.2 | 1.091 |
| EPI_ISL_437451 | hCoV-19/India/GBRC22/2020 | Asia / India / Gujarat / Ahmedabad | 2020-04-26 | D614G | -149.3 | 1.037 |
| EPI_ISL_437452 | hCoV-19/India/GBRC23/2020 | Asia / India / Gujarat / Ahmedabad | 2020-04-26 | D614G:L54F | -137.6 | 1.061 |
| EPI_ISL_437453 | hCoV-19/India/GBRC24a/2020 | Asia / India / Gujarat / Ahmedabad | 2020-04-26 | Ancestral | -129.8 | 1.124 |
| EPI_ISL_437454 | hCoV-19/India/GBRC24b/2020 | Asia / India / Gujarat / Ahmedabad | 2020-04-26 | E583D | -141 | 6.485 |
| EPI_ISL_437537 | hCoV-19/India/S8/2020 | Asia / India / West Bengal / Howrah | 2020-03-30 | D614G | -149.3 | 1.037 |

| | | | | | | |
|---|---|---|---|---|---|---|
| EPI_ISL_437539 | hCoV-19/India/S12/2020 | Asia / India / West Bengal / East Medinipur | 2020-04-06 | D614G | -149.3 | 1.037 |
| EPI_ISL_437626 | hCoV-19/India/GMC-KP1125/2020 | Asia / India / Telangana / Hyderabad | 2020-03-24 | D614G | -149.3 | 1.037 |
| EPI_ISL_438138 | hCoV-19/India/GMC-RR1191/2020 | Asia / India / Telangana / Hyderabad | 2020-03-25 | Ancestral | -129.8 | 1.124 |
| EPI_ISL_438139 | hCoV-19/India/GMC-RK1090/2020 | Asia / India / Telangana / Hyderabad | 2020-03-25 | Ancestral | -129.8 | 1.124 |
| EPI_ISL_444456 | hCoV-19/India/GBRC25/2020 | Asia / India / Gujarat / Ahmedabad | 2020-04-26 | D614G | -149.3 | 1.037 |
| EPI_ISL_444457 | hCoV-19/India/GBRC26/2020 | Asia / India / Gujarat / Ahmedabad | 2020-04-26 | D614G | -149.3 | 1.037 |
| EPI_ISL_444458 | hCoV-19/India/GBRC27/2020 | Asia / India / Gujarat / Ahmedabad | 2020-04-26 | D614G | -149.3 | 1.037 |
| EPI_ISL_444459 | hCoV-19/India/GBRC28/2020 | Asia / India / Gujarat / Ahmedabad | 2020-04-30 | D614G:E583D | -128.8 | 1.091 |
| EPI_ISL_444460 | hCoV-19/India/GBRC29/2020 | Asia / India / Gujarat / Ahmedabad | 2020-04-30 | D614G | -149.3 | 1.037 |
| EPI_ISL_444461 | hCoV-19/India/GBRC30/2020 | Asia / India / Gujarat / Ahmedabad | 2020-04-29 | D614G | -149.3 | 1.037 |

| | | | | | | |
|---|---|---|---|---|---|---|
| EPI_ISL_444462 | hCoV-19/India/GBRC31/2020 | Asia / India / Gujarat / Ahmedabad | 2020-04-29 | D614G | -149.3 | 1.037 |
| EPI_ISL_444463 | hCoV-19/India/GBRC32/2020 | Asia / India / Gujarat / Ahmedabad | 2020-04-29 | D614G:Q677H | -145.2 | 1.091 |
| EPI_ISL_444464 | hCoV-19/India/GBRC33/2020 | Asia / India / Gujarat / Ahmedabad | 2020-04-29 | D614G | -149.3 | 1.037 |
| EPI_ISL_444465 | hCoV-19/India/GBRC34/2020 | Asia / India / Gujarat / Ahmedabad | 2020-04-29 | D614G:T572I | -145.6 | 8.737 |
| EPI_ISL_444466 | hCoV-19/India/GBRC35/2020 | Asia / India / Gujarat / Ahmedabad | 2020-04-29 | D614G:T572I | -145.6 | 8.737 |
| EPI_ISL_444467 | hCoV-19/India/GBRC36/2020 | Asia / India / Gujarat / Ahmedabad | 2020-04-29 | D614G | -149.3 | 1.037 |
| EPI_ISL_444468 | hCoV-19/India/GBRC37/2020 | Asia / India / Gujarat / Ahmedabad | 2020-04-29 | D614G | -149.3 | 1.037 |
| EPI_ISL_444469 | hCoV-19/India/GBRC38/2020 | Asia / India / Gujarat / Ahmedabad | 2020-04-29 | D614G:E583D | -128.8 | 1.091 |
| EPI_ISL_444470 | hCoV-19/India/GBRC39/2020 | Asia / India / Gujarat / Ahmedabad | 2020-04-29 | D614G | -149.3 | 1.037 |
| EPI_ISL_444471 | hCoV-19/India/GBRC40/2020 | Asia / India / Gujarat / Ahmedabad | 2020-04-29 | D614G | -149.3 | 1.037 |

| | | | | | | |
|---|---|---|---|---|---|---|
| EPI_ISL_444472 | hCoV-19/India/GBRC41/2020 | Asia / India / Gujarat / Ahmedabad | 2020-04-29 | D614G | -149.3 | 1.037 |
| EPI_ISL_444473 | hCoV-19/India/GBRC42/2020 | Asia / India / Gujarat / Ahmedabad | 2020-04-29 | D614G:A892V | -143 | 3.897 |
| EPI_ISL_444474 | hCoV-19/India/GBRC43/2020 | Asia / India / Gujarat / Ahmedabad | 2020-04-29 | D614G:T827I | -124.5 | 5.789 |
| EPI_ISL_444475 | hCoV-19/India/GBRC44/2020 | Asia / India / Gujarat / Ahmedabad | 2020-04-29 | D614G | -149.3 | 1.037 |
| EPI_ISL_444476 | hCoV-19/India/GBRC45/2020 | Asia / India / Gujarat / Ahmedabad | 2020-04-29 | D614G:T572I | -145.6 | 8.737 |
| EPI_ISL_444477 | hCoV-19/India/GBRC46/2020 | Asia / India / Gujarat / Ahmedabad | 2020-04-29 | D614G | -149.3 | 1.037 |
| EPI_ISL_444478 | hCoV-19/India/GBRC47/2020 | Asia / India / Gujarat / Ahmedabad | 2020-04-30 | D614G:E156D | -146.6 | 2.590 |
| EPI_ISL_444479 | hCoV-19/India/GBRC48/2020 | Asia / India / Gujarat / Ahmedabad | 2020-04-29 | D614G | -149.3 | 1.037 |
| EPI_ISL_444480 | hCoV-19/India/GBRC49/2020 | Asia / India / Gujarat / Ahmedabad | 2020-04-30 | D614G | -149.3 | 1.037 |
| EPI_ISL_444481 | hCoV-19/India/GBRC50/2020 | Asia / India / Gujarat / Ahmedabad | 2020-04-30 | D614G | -149.3 | 1.037 |

| | | | | | | |
|---|---|---|---|---|---|---|
| EPI_ISL_444482 | hCoV-19/India/GBRC51/2020 | Asia / India / Gujarat / Prantij | 2020-05-02 | D614G | -149.3 | 1.037 |
| EPI_ISL_444483 | hCoV-19/India/GBRC52/2020 | Asia / India / Gujarat / Prantij | 2020-05-02 | D614G | -149.3 | 1.037 |
| EPI_ISL_444484 | hCoV-19/India/GBRC53/2020 | Asia / India / Gujarat / Modasa | 2020-05-04 | D614G:R78M | -147.1 | 4.033 |
| EPI_ISL_444485 | hCoV-19/India/GBRC54/2020 | Asia / India / Gujarat / Modasa | 2020-05-04 | D614G:R78M | -147.1 | 4.033 |
| EPI_ISL_444486 | hCoV-19/India/GBRC55/2020 | Asia / India / Gujarat / Dhansura | 2020-05-04 | D614G:L54F | -137.6 | 1.061 |
| EPI_ISL_447030 | hCoV-19/India/GBRC56a/2020 | Asia / India / Gujarat / Ahmedabad | 2020-05-03 | D614G:S13I | NA | NA |
| EPI_ISL_447031 | hCoV-19/India/GBRC56b/2020 | Asia / India / Gujarat / Ahmedabad | 2020-05-03 | D614G | -149.3 | 1.037 |
| EPI_ISL_447033 | hCoV-19/India/GBRC58/2020 | Asia / India / Gujarat / Ahmedabad | 2020-05-03 | D614G:L54F | -137.6 | 1.061 |
| EPI_ISL_447035 | hCoV-19/India/GBRC60/2020 | Asia / India / Gujarat / Ahmedabad | 2020-05-03 | D614G:T572I | -145.6 | 8.737 |
| EPI_ISL_447036 | hCoV-19/India/GBRC61/2020 | Asia / India / Gujarat / Ahmedabad | 2020-05-03 | D614G | -149.3 | 1.037 |

| | | | | | | |
|---|---|---|---|---|---|---|
| EPI_ISL_447037 | hCoV-19/India/GBRC62/2020 | Asia / India / Gujarat / Ahmedabad | 2020-05-03 | D614G:V1104L | -152.1 | 1.600 |
| EPI_ISL_447038 | hCoV-19/India/GBRC63/2020 | Asia / India / Gujarat / Ahmedabad | 2020-05-03 | D614G:F2L | NA | NA |
| EPI_ISL_447039 | hCoV-19/India/GBRC64/2020 | Asia / India / Gujarat / Ahmedabad | 2020-05-03 | D614G | -149.3 | 1.037 |
| EPI_ISL_447040 | hCoV-19/India/GBRC65a/2020 | Asia / India / Gujarat / Ahmedabad | 2020-05-03 | D614G:M177I | -156.3 | 0.917 |
| EPI_ISL_447041 | hCoV-19/India/GBRC65b/2020 | Asia / India / Gujarat / Ahmedabad | 2020-05-03 | D614G:M177I | -156.3 | 0.917 |
| EPI_ISL_447042 | hCoV-19/India/GBRC66/2020 | Asia / India / Gujarat / Ahmedabad | 2020-05-03 | D614G:A879S | -146.6 | 21.300 |
| EPI_ISL_447043 | hCoV-19/India/GBRC67a/2020 | Asia / India / Gujarat / Ahmedabad | 2020-05-03 | D614G:E583D | -128.8 | 1.091 |
| EPI_ISL_447044 | hCoV-19/India/GBRC67b/2020 | Asia / India / Gujarat / Ahmedabad | 2020-05-03 | D614G:E583D | -128.8 | 1.091 |
| EPI_ISL_447045 | hCoV-19/India/GBRC68/2020 | Asia / India / Gujarat / Ahmedabad | 2020-05-03 | D614G | -149.3 | 1.037 |
| EPI_ISL_447046 | hCoV-19/India/GBRC69/2020 | Asia / India / Gujarat / Ahmedabad | 2020-05-04 | D614G:L54F:E471Q | -144.3 | 2.941 |

| | | | | | | |
|---|---|---|---|---|---|---|
| EPI_ISL_447047 | hCoV-19/India/GBRC70/2020 | Asia / India / Gujarat / Gandhinagar | 2020-04-29 | D614G | -149.3 | 1.037 |
| EPI_ISL_447048 | hCoV-19/India/GBRC71/2020 | Asia / India / Gujarat / Gandhinagar | 2020-04-27 | D614G | -149.3 | 1.037 |
| EPI_ISL_447049 | hCoV-19/India/GBRC72a/2020 | Asia / India / Gujarat / Gandhinagar | 2020-04-29 | D614G:A706S | -136 | 4.137 |
| EPI_ISL_447050 | hCoV-19/India/GBRC72b/2020 | Asia / India / Gujarat / Gandhinagar | 2020-04-29 | D614G | -149.3 | 1.037 |
| EPI_ISL_447051 | hCoV-19/India/GBRC73/2020 | Asia / India / Gujarat / Gandhinagar | 2020-04-29 | D614G | -149.3 | 1.037 |
| EPI_ISL_447052 | hCoV-19/India/GBRC74/2020 | Asia / India / Gujarat / Gandhinagar | 2020-05-02 | D614G | -149.3 | 1.037 |
| EPI_ISL_447053 | hCoV-19/India/GBRC75/2020 | Asia / India / Gujarat / Gandhinagar | 2020-04-29 | D614G | -149.3 | 1.037 |
| EPI_ISL_447534 | hCoV-19/India/GBRC76/2020 | Asia / India / Gujarat / Modasa | 2020-05-05 | D614G:R78M | -147.1 | 4.033 |
| EPI_ISL_447535 | hCoV-19/India/GBRC77/2020 | Asia / India / Gujarat / Modasa | 2020-05-05 | D614G:R78M | -147.1 | 4.033 |
| EPI_ISL_447536 | hCoV-19/India/GBRC78a/2020 | Asia / India / Gujarat / Modasa | 2020-05-05 | D614G | -149.3 | 1.037 |

| | | | | | | |
|---|---|---|---|---|---|---|
| EPI_ISL_447537 | hCoV-19/India/GBRC78b/2020 | Asia / India / Gujarat / Modasa | 2020-05-05 | D614G:R78M | -147.1 | 4.033 |
| EPI_ISL_447538 | hCoV-19/India/GBRC79a/2020 | Asia / India / Gujarat / Modasa | 2020-05-05 | D614G | -149.3 | 1.037 |
| EPI_ISL_447539 | hCoV-19/India/GBRC79b/2020 | Asia / India / Gujarat / Modasa | 2020-05-05 | D614G | -149.3 | 1.037 |
| EPI_ISL_447540 | hCoV-19/India/GBRC81a/2020 | Asia / India / Gujarat / Modasa | 2020-05-05 | D614G:R78M | -147.1 | 4.033 |
| EPI_ISL_447541 | hCoV-19/India/GBRC81b/2020 | Asia / India / Gujarat / Modasa | 2020-05-05 | D614G:R78M | -147.1 | 4.033 |
| EPI_ISL_447542 | hCoV-19/India/GBRC82a/2020 | Asia / India / Gujarat / Modasa | 2020-05-05 | D614G:R78M | -147.1 | 4.033 |
| EPI_ISL_447543 | hCoV-19/India/GBRC82b/2020 | Asia / India / Gujarat / Modasa | 2020-05-05 | D614G:R78M | -147.1 | 4.033 |
| EPI_ISL_447544 | hCoV-19/India/GBRC83/2020 | Asia / India / Gujarat / Modasa | 2020-05-05 | D614G | -149.3 | 1.037 |
| EPI_ISL_447545 | hCoV-19/India/GBRC84/2020 | Asia / India / Gujarat / Prantij | 2020-05-05 | D614G | -149.3 | 1.037 |
| EPI_ISL_447546 | hCoV-19/India/GBRC85/2020 | Asia / India / Gujarat / Himatnagar | 2020-05-05 | D614G:E583D | -128.8 | 1.091 |

| | | | | | | |
|---|---|---|---|---|---|---|
| EPI_ISL_447547 | hCoV-19/India/GBRC86/2020 | Asia / India / Gujarat / Gandhinagar | 2020-04-28 | D614G | -149.3 | 1.037 |
| EPI_ISL_447548 | hCoV-19/India/GBRC87/2020 | Asia / India / Gujarat / Dahegam | 2020-04-28 | D614G | -149.3 | 1.037 |
| EPI_ISL_447549 | hCoV-19/India/GBRC88a/2020 | Asia / India / Gujarat / Dahegam | 2020-05-03 | D614G | -149.3 | 1.037 |
| EPI_ISL_447550 | hCoV-19/India/GBRC88b/2020 | Asia / India / Gujarat / Dahegam | 2020-05-03 | D614G | -149.3 | 1.037 |
| EPI_ISL_447551 | hCoV-19/India/GBRC89/2020 | Asia / India / Gujarat / Gandhinagar | 2020-04-25 | D614G | -149.3 | 1.037 |
| EPI_ISL_447552 | hCoV-19/India/GBRC90/2020 | Asia / India / Gujarat / Gandhinagar | 2020-05-02 | D614G | -149.3 | 1.037 |
| EPI_ISL_447553 | hCoV-19/India/GBRC91/2020 | Asia / India / Gujarat / Gandhinagar | 2020-04-27 | D614G | -149.3 | 1.037 |
| EPI_ISL_447554 | hCoV-19/India/GBRC92/2020 | Asia / India / Gujarat / Gandhinagar | 2020-04-26 | D614G | -149.3 | 1.037 |
| EPI_ISL_447555 | hCoV-19/India/GBRC93/2020 | Asia / India / Gujarat / Mansa | 2020-04-28 | D614G | -149.3 | 1.037 |
| EPI_ISL_447556 | hCoV-19/India/CCMB_J021/2020 | Asia / India / Telangana | 2020-03-30 | K77M | -84.8 | 0.194 |
| EPI_ISL_447557 | hCoV-19/India/CCMB_J009/2020 | Asia / India / Telangana | 2020-03-30 | Ancestral | -129.8 | 1.124 |

| | | | | | | |
|---|---|---|---|---|---|---|
| EPI_ISL_447558 | hCoV-19/India/CCMB_J014/2020 | Asia / India / Telangana | 2020-03-30 | Ancestral | -129.8 | 1.124 |
| EPI_ISL_447559 | hCoV-19/India/CCMB_J048/2020 | Asia / India / Telangana | 2020-03-31 | Ancestral | -129.8 | 1.124 |
| EPI_ISL_447560 | hCoV-19/India/CCMB_J043/2020 | Asia / India / Telangana | 2020-03-31 | Ancestral | -129.8 | 1.124 |
| EPI_ISL_447561 | hCoV-19/India/CCMB_J067/2020 | Asia / India / Telangana | 2020-03-31 | Ancestral | -129.8 | 1.124 |
| EPI_ISL_447562 | hCoV-19/India/CCMB_J068/2020 | Asia / India / Telangana | 2020-03-31 | Ancestral | -129.8 | 1.124 |
| EPI_ISL_447563 | hCoV-19/India/CCMB_J057/2020 | Asia / India / Telangana | 2020-03-31 | Ancestral | -129.8 | 1.124 |
| EPI_ISL_447564 | hCoV-19/India/CCMB_J212/2020 | Asia / India / Telangana | 2020-04-01 | Ancestral | -129.8 | 1.124 |
| EPI_ISL_447565 | hCoV-19/India/CCMB_J278/2020 | Asia / India / Telangana | 2020-04-01 | Ancestral | -129.8 | 1.124 |
| EPI_ISL_447566 | hCoV-19/India/CCMB_J131/2020 | Asia / India / Telangana | 2020-04-01 | K77M | -84.8 | 0.194 |
| EPI_ISL_447567 | hCoV-19/India/CCMB_J304/2020 | Asia / India / Telangana | 2020-04-01 | Ancestral | -129.8 | 1.124 |
| EPI_ISL_447568 | hCoV-19/India/CCMB_J166/2020 | Asia / India / Telangana | 2020-04-01 | Ancestral | -129.8 | 1.124 |
| EPI_ISL_447569 | hCoV-19/India/CCMB_J206/2020 | Asia / India / Telangana | 2020-04-01 | Ancestral | -129.8 | 1.124 |
| EPI_ISL_447570 | hCoV-19/India/CCMB_J207/2020 | Asia / India / Telangana | 2020-04-01 | Ancestral | -129.8 | 1.124 |
| EPI_ISL_447571 | hCoV-19/India/CCMB_J224/2020 | Asia / India / Telangana | 2020-04-01 | K77M:S1021F | -128.5 | 0.354 |

| | | | | | | |
|---|---|---|---|---|---|---|
| EPI_ISL_447572 | hCoV-19/India/CCMB_J230/2020 | Asia / India / Telangana | 2020-04-01 | Ancestral | -129.8 | 1.124 |
| EPI_ISL_447573 | hCoV-19/India/CCMB_J118/2020 | Asia / India / Telangana | 2020-04-01 | Ancestral | -129.8 | 1.124 |
| EPI_ISL_447574 | hCoV-19/India/CCMB_J208/2020 | Asia / India / Telangana | 2020-04-01 | Ancestral | -129.8 | 1.124 |
| EPI_ISL_447575 | hCoV-19/India/CCMB_J458/2020 | Asia / India / Telangana | 2020-04-02 | Ancestral | -129.8 | 1.124 |
| EPI_ISL_447576 | hCoV-19/India/CCMB_J232/2020 | Asia / India / Telangana | 2020-04-02 | Ancestral | -129.8 | 1.124 |
| EPI_ISL_447577 | hCoV-19/India/CCMB_J321/2020 | Asia / India / Telangana | 2020-04-02 | Ancestral | -129.8 | 1.124 |
| EPI_ISL_447578 | hCoV-19/India/CCMB_J327/2020 | Asia / India / Telangana | 2020-04-02 | Ancestral | -129.8 | 1.124 |
| EPI_ISL_447579 | hCoV-19/India/CCMB_K128/2020 | Asia / India / Telangana | 2020-04-10 | Ancestral | -129.8 | 1.124 |
| EPI_ISL_447580 | hCoV-19/India/CCMB_K496/2020 | Asia / India / Telangana | 2020-04-13 | Ancestral | -129.8 | 1.124 |
| EPI_ISL_447581 | hCoV-19/India/CCMB_K600B/2020 | Asia / India / Telangana | 2020-04-14 | Ancestral | -129.8 | 1.124 |
| EPI_ISL_447582 | hCoV-19/India/CCMB_K601/2020 | Asia / India / Telangana | 2020-04-14 | Ancestral | -129.8 | 1.124 |
| EPI_ISL_447583 | hCoV-19/India/CCMB_L007/2020 | Asia / India / Telangana | 2020-04-16 | Ancestral | -129.8 | 1.124 |

| | | | | | | |
|---|---|---|---|---|---|---|
| EPI_ISL_447584 | hCoV-19/India/CCMB_C1-12/2020 | Asia / India / Tamil Nadu | 2020-04-16 | Ancestral | -129.8 | 1.124 |
| EPI_ISL_447585 | hCoV-19/India/CCMB_C2-13/2020 | Asia / India / Tamil Nadu | 2020-04-16 | Ancestral | -129.8 | 1.124 |
| EPI_ISL_447586 | hCoV-19/India/CCMB_C3-14/2020 | Asia / India / Tamil Nadu | 2020-04-16 | Ancestral | -129.8 | 1.124 |
| EPI_ISL_447587 | hCoV-19/India/CCMB_C4-15/2020 | Asia / India / Tamil Nadu | 2020-04-20 | D614G | -149.3 | 1.037 |
| EPI_ISL_447847 | hCoV-19/India/CCMB_J045/2020 | Asia / India / Telangana | 2020-03-31 | Ancestral | -129.8 | 1.124 |
| EPI_ISL_447848 | hCoV-19/India/CCMB_J165/2020 | Asia / India / Telangana | 2020-04-01 | Ancestral | -129.8 | 1.124 |
| EPI_ISL_447849 | hCoV-19/India/CCMB_J107/2020 | Asia / India / Telangana | 2020-04-01 | Ancestral | -129.8 | 1.124 |
| EPI_ISL_447850 | hCoV-19/India/CCMB_J132/2020 | Asia / India / Telangana | 2020-04-01 | K77M | -84.8 | 0.194 |
| EPI_ISL_447851 | hCoV-19/India/CCMB_J223/2020 | Asia / India / Telangana | 2020-04-01 | Ancestral | -129.8 | 1.124 |
| EPI_ISL_447852 | hCoV-19/India/CCMB_J235/2020 | Asia / India / Telangana | 2020-04-01 | Ancestral | -129.8 | 1.124 |
| EPI_ISL_447853 | hCoV-19/India/CCMB_J247/2020 | Asia / India / Telangana | 2020-04-01 | Ancestral | -129.8 | 1.124 |
| EPI_ISL_447854 | hCoV-19/India/CCMB_J350/2020 | Asia / India / Telangana | 2020-04-02 | Ancestral | -129.8 | 1.124 |
| EPI_ISL_447855 | hCoV-19/India/CCMB_J318/2020 | Asia / India / Telangana | 2020-04-02 | Ancestral | -129.8 | 1.124 |
| EPI_ISL_447856 | hCoV-19/India/CCMB_J374/2020 | Asia / India / Telangana | 2020-04-02 | K77M | -84.8 | 0.194 |

| | | | | | | |
|---|---|---|---|---|---|---|
| EPI_ISL_447857 | hCoV-19/India/CCMB_J390/2020 | Asia / India / Telangana | 2020-04-02 | Ancestral | -129.8 | 1.124 |
| EPI_ISL_447858 | hCoV-19/India/CCMB_J812/2020 | Asia / India / Telangana | 2020-04-06 | K77M | -84.8 | 0.194 |
| EPI_ISL_447859 | hCoV-19/India/CCMB_K771/2020 | Asia / India / Telangana | 2020-04-15 | Ancestral | -129.8 | 1.124 |
| EPI_ISL_447860 | hCoV-19/India/CCMB_K849/2020 | Asia / India / Telangana | 2020-04-16 | Ancestral | -129.8 | 1.124 |
| EPI_ISL_447861 | hCoV-19/India/CCMB_L149/2020 | Asia / India / Telangana | 2020-04-16 | Ancestral | -129.8 | 1.124 |
| EPI_ISL_447862 | hCoV-19/India/CCMB_J047/2020 | Asia / India / Telangana | 2020-03-31 | Ancestral | -129.8 | 1.124 |
| EPI_ISL_447863 | hCoV-19/India/CCMB_K599/2020 | Asia / India / Telangana | 2020-04-14 | Ancestral | -129.8 | 1.124 |
| EPI_ISL_447864 | hCoV-19/India/CCMB_L044/2020 | Asia / India / Telangana | 2020-04-16 | Ancestral | -129.8 | 1.124 |
| EPI_ISL_447865 | hCoV-19/India/CCMB_K499/2020 | Asia / India / Telangana | 2020-04-14 | Ancestral | -129.8 | 1.124 |
| EPI_ISL_447866 | hCoV-19/India/CCMB_K500/2020 | Asia / India / Telangana | 2020-04-14 | T602I | -122.2 | 1.061 |
| EPI_ISL_450321 | hCoV-19/India/CCMB_NIV1/2020 | Asia / India / Maharashtra | 2020-04-04 | Ancestral | -129.8 | 1.124 |

| | | | | | | |
|---|---|---|---|---|---|---|
| EPI_ISL_450322 | hCoV-19/India/CCMB_NIV2/2020 | Asia / India / Maharashtra | 2020-04-04 | Ancestral | -129.8 | 1.124 |
| EPI_ISL_450323 | hCoV-19/India/CCMB_NIV3/2020 | Asia / India / Maharashtra | 2020-03-27 | D614G | -149.3 | 1.037 |
| EPI_ISL_450324 | hCoV-19/India/CCMB_NIV4/2020 | Asia / India / Maharashtra | 2020-03-26 | Ancestral | -129.8 | 1.124 |
| EPI_ISL_450325 | hCoV-19/India/CCMB_NIV5/2020 | Asia / India / Maharashtra | 2020-03-17 | D614G | -149.3 | 1.037 |
| EPI_ISL_450326 | hCoV-19/India/CCMB_J125/2020 | Asia / India / Telangana | 2020-04-01 | Ancestral | -129.8 | 1.124 |
| EPI_ISL_450327 | hCoV-19/India/CCMB_J199/2020 | Asia / India / Telangana | 2020-04-01 | Ancestral | -129.8 | 1.124 |
| EPI_ISL_450328 | hCoV-19/India/CCMB_J283/2020 | Asia / India / Telangana | 2020-04-01 | Ancestral | -129.8 | 1.124 |
| EPI_ISL_450329 | hCoV-19/India/CCMB_J288/2020 | Asia / India / Telangana | 2020-04-01 | Ancestral | -129.8 | 1.124 |
| EPI_ISL_450330 | hCoV-19/India/CCMB_J375/2020 | Asia / India / Telangana | 2020-04-02 | K77M | -84.8 | 0.194 |
| EPI_ISL_450331 | hCoV-19/India/CCMB_J220/2020 | Asia / India / Telangana | 2020-04-01 | Ancestral | -129.8 | 1.124 |
| EPI_ISL_450332 | hCoV-19/India/CCMB_J222/2020 | Asia / India / Telangana | 2020-04-01 | Ancestral | -129.8 | 1.124 |
| EPI_ISL_450781 | hCoV-19/India/GBRC94/2020 | Asia / India / Gujarat / Junagadh | 2020-05-09 | D614G | -149.3 | 1.037 |

| | | | | | | |
|---|---|---|---|---|---|---|
| EPI_ISL_450782 | hCoV-19/India/GBRC95a/2020 | Asia / India / Gujarat / Ahmedabad | 2020-04-11 | D614G:H1083Q:E583D | -152.4 | 3.817 |
| EPI_ISL_450783 | hCoV-19/India/GBRC95b/2020 | Asia / India / Gujarat / Ahmedabad | 2020-04-11 | D614G:H1083Q:R78M | -141.7 | 1.131 |
| EPI_ISL_450784 | hCoV-19/India/GBRC96/2020 | Asia / India / Gujarat / Botad | 2020-05-09 | Ancestral | -129.8 | 1.124 |
| EPI_ISL_450785 | hCoV-19/India/GBRC97a/2020 | Asia / India / Gujarat / Una | 2020-05-10 | D614G | -149.3 | 1.037 |
| EPI_ISL_450786 | hCoV-19/India/GBRC97b/2020 | Asia / India / Gujarat / Una | 2020-05-10 | D614G | -149.3 | 1.037 |
| EPI_ISL_450787 | hCoV-19/India/GBRC98/2020 | Asia / India / Gujarat / Una | 2020-05-11 | Q677H | -118.9 | 3.498 |
| EPI_ISL_450788 | hCoV-19/India/GBRC99/2020 | Asia / India / Gujarat / Rajkot | 2020-05-06 | D614G | -149.3 | 1.037 |
| EPI_ISL_450789 | hCoV-19/India/GBRC100/2020 | Asia / India / Gujarat / Kodinar | 2020-05-10 | D614G | -149.3 | 1.037 |
| EPI_ISL_450790 | hCoV-19/India/GBRC101/2020 | Asia / India / Gujarat / Rajkot | 2020-04-28 | Ancestral | -129.8 | 1.124 |
| EPI_ISL_450791 | hCoV-19/India/GBRC102/2020 | Asia / India / Gujarat / Rajkot | 2020-04-27 | Ancestral | -129.8 | 1.124 |
| EPI_ISL_451149 | hCoV-19/India/GBRC104/2020 | Asia / India / Gujarat / Jamnagar | 2020-05-07 | D614G | -149.3 | 1.037 |

| | | | | | | |
|---|---|---|---|---|---|---|
| EPI_ISL_451150 | hCoV-19/India/GBRC105/2020 | Asia / India / Gujarat / Jamnagar | 2020-05-06 | D614G:L5F:S162I | NA | NA |
| EPI_ISL_451151 | hCoV-19/India/GBRC106/2020 | Asia / India / Gujarat / Jamnagar | 2020-05-05 | D614G | -149.3 | 1.037 |
| EPI_ISL_451152 | hCoV-19/India/GBRC107/2020 | Asia / India / Gujarat / Jamnagar | 2020-05-09 | D614G | -149.3 | 1.037 |
| EPI_ISL_451153 | hCoV-19/India/GBRC108/2020 | Asia / India / Gujarat / Jamnagar | 2020-05-10 | D614G:L54F | -137.6 | 1.061 |
| EPI_ISL_451154 | hCoV-19/India/GBRC109/2020 | Asia / India / Gujarat / Dahod | 2020-05-03 | Ancestral | -129.8 | 1.124 |
| EPI_ISL_451155 | hCoV-19/India/GBRC110/2020 | Asia / India / Gujarat / Vadodara | 2020-05-03 | D614G | -149.3 | 1.037 |
| EPI_ISL_451156 | hCoV-19/India/GBRC111/2020 | Asia / India / Gujarat / Dahod | 2020-05-01 | Ancestral | -129.8 | 1.124 |
| EPI_ISL_451157 | hCoV-19/India/GBRC112/2020 | Asia / India / Gujarat / Vadodara | 2020-05-02 | D614G | -149.3 | 1.037 |
| EPI_ISL_451158 | hCoV-19/India/GBRC113/2020 | Asia / India / Gujarat / Vadodara | 2020-05-03 | D614G | -149.3 | 1.037 |
| EPI_ISL_451159 | hCoV-19/India/GBRC114/2020 | Asia / India / Gujarat / Dahod | 2020-05-03 | Ancestral | -129.8 | 1.124 |

| | | | | | | |
|---|---|---|---|---|---|---|
| EPI_ISL_451160 | hCoV-19/India/GBRC115/2020 | Asia / India / Gujarat / Vadodara | 2020-05-03 | D614G | -149.3 | 1.037 |
| EPI_ISL_451161 | hCoV-19/India/GBRC116/2020 | Asia / India / Gujarat / Dahod | 2020-05-03 | Ancestral | -129.8 | 1.124 |
| EPI_ISL_451162 | hCoV-19/India/GBRC117/2020 | Asia / India / Gujarat / Vadodara | 2020-05-03 | D614G | -149.3 | 1.037 |
| EPI_ISL_451163 | hCoV-19/India/GBRC118/2020 | Asia / India / Gujarat / Vadodara | 2020-05-02 | D614G | -149.3 | 1.037 |
| EPI_ISL_451666 | hCoV-19/India/GBRC103/2020 | Asia / India / Gujarat / Jamnagar | 2020-05-04 | Ancestral | -129.8 | 1.124 |
| EPI_ISL_452192 | hCoV-19/India/NIV-10380/2020 | Asia / India / Maharashtra | 2020-04-16 | D614G | -149.3 | 1.037 |
| EPI_ISL_452193 | hCoV-19/India/NIV-10419/2020 | Asia / India / Maharashtra | 2020-04-16 | D614G | -149.3 | 1.037 |
| EPI_ISL_452194 | hCoV-19/India/NIV-11677/2020 | Asia / India / Maharashtra | 2020-04-19 | D614G | -149.3 | 1.037 |
| EPI_ISL_452195 | hCoV-19/India/NIV-11681/2020 | Asia / India / Maharashtra | 2020-04-19 | D614G:T22I | NA | NA |
| EPI_ISL_452196 | hCoV-19/India/NIV-11683/2020 | Asia / India / Maharashtra | 2020-04-19 | D614G:T22I | NA | NA |
| EPI_ISL_452197 | hCoV-19/India/NIV-11687/2020 | Asia / India / Maharashtra | 2020-04-19 | D614G:Q677H | -145.2 | 1.091 |
| EPI_ISL_452198 | hCoV-19/India/NIV-11855/2020 | Asia / India / Maharashtra | 2020-04-20 | D614G | -149.3 | 1.037 |

| | | | | | | |
|---|---|---|---|---|---|---|
| EPI_ISL_452199 | hCoV-19/India/NIV-12019/2020 | Asia / India / Maharashtra | 2020-04-20 | D614G | -149.3 | 1.037 |
| EPI_ISL_452200 | hCoV-19/India/NIV-12024/2020 | Asia / India / Maharashtra | 2020-04-20 | D614G | -149.3 | 1.037 |
| EPI_ISL_452201 | hCoV-19/India/NIV-12067/2020 | Asia / India / Maharashtra | 2020-04-20 | D614G | -149.3 | 1.037 |
| EPI_ISL_452202 | hCoV-19/India/NIV-2810/2020 | Asia / India / Maharashtra | 2020-03-16 | Ancestral | -129.8 | 1.124 |
| EPI_ISL_452203 | hCoV-19/India/NIV-4367/2020 | Asia / India / Maharashtra | 2020-03-22 | Ancestral | -129.8 | 1.124 |
| EPI_ISL_452204 | hCoV-19/India/NIV-5109/2020 | Asia / India / Maharashtra | 2020-03-26 | Ancestral | -129.8 | 1.124 |
| EPI_ISL_452205 | hCoV-19/India/NIV-5119/2020 | Asia / India / Maharashtra | 2020-03-26 | Ancestral | -129.8 | 1.124 |
| EPI_ISL_452206 | hCoV-19/India/NIV-6311/2020 | Asia / India / Maharashtra | 2020-03-30 | Ancestral | -129.8 | 1.124 |
| EPI_ISL_452207 | hCoV-19/India/NIV-7439/2020 | Asia / India / Maharashtra | 2020-04-05 | D614G | -149.3 | 1.037 |
| EPI_ISL_452208 | hCoV-19/India/NIV-7511/2020 | Asia / India / Maharashtra | 2020-04-05 | Ancestral | -129.8 | 1.124 |
| EPI_ISL_452209 | hCoV-19/India/NIV-7786/2020 | Asia / India / Maharashtra | 2020-04-06 | Ancestral | -129.8 | 1.124 |
| EPI_ISL_452210 | hCoV-19/India/NIV-7830/2020 | Asia / India / Maharashtra | 2020-04-06 | D614G:Q677H:K558N | -133.8 | 2.711 |
| EPI_ISL_452211 | hCoV-19/India/NIV-9157/2020 | Asia / India / Maharashtra | 2020-04-13 | D614G | -149.3 | 1.037 |
| EPI_ISL_452212 | hCoV-19/India/NIV-9768/2020 | Asia / India / Maharashtra | 2020-04-15 | D614G:F797C | -147.5 | 1.147 |

| | | | | | | |
|---|---|---|---|---|---|---|
| EPI_ISL_452213 | hCoV-19/India/NIV-981/2020 | Asia / India / Maharashtra | 2020-03-10 | Ancestral | -129.8 | 1.124 |
| EPI_ISL_452214 | hCoV-19/India/NIV-QA-709/2020 | Asia / India / Maharashtra | 2020-04-07 | D614G | -149.3 | 1.037 |
| EPI_ISL_452215 | hCoV-19/India/NIV-QC-797/2020 | Asia / India / Maharashtra | 2020-04-22 | Q677H | -118.9 | 3.498 |
| EPI_ISL_452216 | hCoV-19/India/NIV-QC-802/2020 | Asia / India / Maharashtra | 2020-04-27 | D614G | -149.3 | 1.037 |
| EPI_ISL_452217 | hCoV-19/India/NIV-QC-800/2020 | Asia / India / Maharashtra | 2020-04-26 | D614G | -149.3 | 1.037 |
| EPI_ISL_452787 | hCoV-19/India/NIHSAD-2004-18/2020 | Asia / India / Madhya Pradesh | 2020-04-19 | D614G | -149.3 | 1.037 |
| EPI_ISL_452790 | hCoV-19/India/NIHSAD-2604-22/2020 | Asia / India / Madhya Pradesh | 2020-04-24 | D614G | -149.3 | 1.037 |
| EPI_ISL_452792 | hCoV-19/India/NIHSAD-0105-77/2020 | Asia / India / Madhya Pradesh | 2020-04-30 | D614G:C1243F | NA | NA |
| EPI_ISL_452793 | hCoV-19/India/NIHSAD-0405-24/2020 | Asia / India / Madhya Pradesh | 2020-05-03 | Ancestral | -129.8 | 1.124 |
| EPI_ISL_452794 | hCoV-19/India/NIHSAD-0905-216/2020 | Asia / India / Madhya Pradesh | 2020-05-08 | D614G | -149.3 | 1.037 |
| EPI_ISL_452795 | hCoV-19/India/NIHSAD-1005-52/2020 | Asia / India / Madhya Pradesh | 2020-05-09 | Ancestral | -129.8 | 1.124 |
| EPI_ISL_454521 | hCoV-19/India/NIV-11182/2020 | Asia / India / Maharashtra | 2020-04-18 | D614G | -149.3 | 1.037 |

| | | | | | | |
|---|---|---|---|---|---|---|
| EPI_ISL_454524 | hCoV-19/India/NIV-12058/2020 | Asia / India / Maharashtra | 2020-04-20 | D614G:P82L | -131.5 | 1.422 |
| EPI_ISL_454525 | hCoV-19/India/NIV-1738/2020 | Asia / India / Maharashtra | 2020-03-12 | Ancestral | -129.8 | 1.124 |
| EPI_ISL_454526 | hCoV-19/India/NIV-1804/2020 | Asia / India / Maharashtra | 2020-03-13 | Ancestral | -129.8 | 1.124 |
| EPI_ISL_454527 | hCoV-19/India/NIV-2387/2020 | Asia / India / Maharashtra | 2020-03-13 | D614G | -149.3 | 1.037 |
| EPI_ISL_454528 | hCoV-19/India/NIV-3472/2020 | Asia / India / Maharashtra | 2020-03-17 | I1179N | NA | NA |
| EPI_ISL_454529 | hCoV-19/India/NIV-4130/2020 | Asia / India / Maharashtra | 2020-03-21 | D614G | -149.3 | 1.037 |
| EPI_ISL_454530 | hCoV-19/India/NIV-4271/2020 | Asia / India / Maharashtra | 2020-03-22 | D614G:E484Q | -150.4 | 0.714 |
| EPI_ISL_454531 | hCoV-19/India/NIV-4369/2020 | Asia / India / Maharashtra | 2020-03-22 | Ancestral | -129.8 | 1.124 |
| EPI_ISL_454532 | hCoV-19/India/NIV-4379/2020 | Asia / India / Maharashtra | 2020-03-23 | D614G | -149.3 | 1.037 |
| EPI_ISL_454533 | hCoV-19/India/NIV-4438/2020 | Asia / India / Maharashtra | 2020-03-23 | D614G | -149.3 | 1.037 |
| EPI_ISL_454534 | hCoV-19/India/NIV-4622/2020 | Asia / India / Maharashtra | 2020-03-24 | Ancestral | -129.8 | 1.124 |
| EPI_ISL_454536 | hCoV-19/India/NIV-5102/2020 | Asia / India / Maharashtra | 2020-03-26 | Ancestral | -129.8 | 1.124 |
| EPI_ISL_454537 | hCoV-19/India/NIV-5104/2020 | Asia / India / Maharashtra | 2020-03-26 | Ancestral | -129.8 | 1.124 |
| EPI_ISL_454539 | hCoV-19/India/NIV-5815-1/2020 | Asia / India / Maharashtra | 2020-04-12 | Ancestral | -129.8 | 1.124 |

| | | | | | | |
|---|---|---|---|---|---|---|
| EPI_ISL_454540 | hCoV-19/India/NIV-6216/2020 | Asia / India / Maharashtra | 2020-04-01 | Ancestral | -129.8 | 1.124 |
| EPI_ISL_454542 | hCoV-19/India/NIV-6315/2020 | Asia / India / Maharashtra | 2020-03-30 | Ancestral | -129.8 | 1.124 |
| EPI_ISL_454543 | hCoV-19/India/NIV-6423/2020 | Asia / India / Maharashtra | 2020-03-31 | Ancestral | -129.8 | 1.124 |
| EPI_ISL_454544 | hCoV-19/India/NIV-6614/2020 | Asia / India / Maharashtra | 2020-04-02 | W258del | -116.7 | 1.637 |
| EPI_ISL_454547 | hCoV-19/India/NIV-7044/2020 | Asia / India / Maharashtra | 2020-04-04 | Ancestral | -129.8 | 1.124 |
| EPI_ISL_454549 | hCoV-19/India/NIV-7558/2020 | Asia / India / Maharashtra | 2020-04-05 | Ancestral | -129.8 | 1.124 |
| EPI_ISL_454551 | hCoV-19/India/NIV-7893/2020 | Asia / India / Maharashtra | 2020-04-07 | Ancestral | -129.8 | 1.124 |
| EPI_ISL_454552 | hCoV-19/India/NIV-7966/2020 | Asia / India / Maharashtra | 2020-04-06 | Ancestral | -129.8 | 1.124 |
| EPI_ISL_454556 | hCoV-19/India/NIV-9127/2020 | Asia / India / Maharashtra | 2020-04-12 | D614G | -149.3 | 1.037 |
| EPI_ISL_454557 | hCoV-19/India/NIV-9154/2020 | Asia / India / Maharashtra | 2020-04-13 | D614G | -149.3 | 1.037 |
| EPI_ISL_454558 | hCoV-19/India/NIV-9314/2020 | Asia / India / Maharashtra | 2020-04-13 | D614G:T22I:L7I | NA | NA |
| EPI_ISL_454560 | hCoV-19/India/NIV-701_3/2020 | Asia / India / Maharashtra | 2020-04-06 | D614G | -149.3 | 1.037 |
| EPI_ISL_454561 | hCoV-19/India/NIV-729_3/2020 | Asia / India / Maharashtra | 2020-04-07 | Ancestral | -129.8 | 1.124 |
| EPI_ISL_454562 | hCoV-19/India/NIV-815_3/2020 | Asia / India / Maharashtra | 2020-04-07 | D614G:R634S | -144.5 | 1.814 |

| | | | | | | |
|---|---|---|---|---|---|---|
| EPI_ISL_454563 | hCoV-19/India/NIV-QA-707/2020 | Asia / India / Maharashtra | 2020-04-10 | D614G:I1179N | NA | NA |
| EPI_ISL_454564 | hCoV-19/India/NIV-QA-708/2020 | Asia / India / Maharashtra | 2020-04-04 | D614G | -149.3 | 1.037 |
| EPI_ISL_454565 | hCoV-19/India/NIV-QA-710/2020 | Asia / India / Maharashtra | 2020-03-31 | D614G | -149.3 | 1.037 |
| EPI_ISL_454566 | hCoV-19/India/NIV-QC-795/2020 | Asia / India / Maharashtra | 2020-04-21 | D614G | -149.3 | 1.037 |
| EPI_ISL_454567 | hCoV-19/India/NIV-QC-796/2020 | Asia / India / Maharashtra | 2020-04-22 | D614G:D574H | -173.1 | 3.875 |
| EPI_ISL_454568 | hCoV-19/India/NIV-QC-798/2020 | Asia / India / Maharashtra | 2020-04-25 | Ancestral | -129.8 | 1.124 |
| EPI_ISL_454569 | hCoV-19/India/NIV-QC-799/2020 | Asia / India / Maharashtra | 2020-04-26 | D614G | -149.3 | 1.037 |
| EPI_ISL_454570 | hCoV-19/India/NIV-QC-801/2020 | Asia / India / Maharashtra | 2020-04-26 | D614G | -149.3 | 1.037 |
| EPI_ISL_454830 | hCoV-19/India/SMSCOV109/2020 | Asia / India / Rajasthan / Jaipur | 2020-04-23 | Ancestral | -129.8 | 1.124 |
| EPI_ISL_454831 | hCoV-19/India/SMSCOV141/2020 | Asia / India / Rajasthan / Jaipur | 2020-04-29 | T1027I | -136.7 | 1.131 |
| EPI_ISL_454832 | hCoV-19/India/SMSCOV161/2020 | Asia / India / Rajasthan / Jaipur | 2020-04-21 | Ancestral | -129.8 | 1.124 |
| EPI_ISL_454833 | hCoV-19/India/SMSCOV175/2020 | Asia / India / Rajasthan / Jaipur | 2020-04-27 | Ancestral | -129.8 | 1.124 |

| | | | | | | |
|---|---|---|---|---|---|---|
| EPI_ISL_454858 | hCoV-19/India/THSTI-BAL_39/2020 | Asia / India / Haryana | 2020-04-07 | Ancestral | -129.8 | 1.124 |
| EPI_ISL_454859 | hCoV-19/India/THSTI-BAL_41/2020 | Asia / India / Haryana | 2020-04-07 | Ancestral | -129.8 | 1.124 |
| EPI_ISL_454860 | hCoV-19/India/THSTI-BAL_42/2020 | Asia / India / Haryana | 2020-04-07 | Ancestral | -129.8 | 1.124 |
| EPI_ISL_454861 | hCoV-19/India/THSTI-BAL_47/2020 | Asia / India / Haryana | 2020-04-07 | Ancestral | -129.8 | 1.124 |
| EPI_ISL_454862 | hCoV-19/India/THSTI-BAL_231/2020 | Asia / India / Haryana | 2020-04-11 | Ancestral | -129.8 | 1.124 |
| EPI_ISL_454863 | hCoV-19/India/THSTI-BAL_360/2020 | Asia / India / Haryana | 2020-04-13 | Ancestral | -129.8 | 1.124 |
| EPI_ISL_454864 | hCoV-19/India/THSTI-BAL_912/2020 | Asia / India / Haryana | 2020-04-16 | Ancestral | -129.8 | 1.124 |
| EPI_ISL_454865 | hCoV-19/India/THSTI-BAL_913/2020 | Asia / India / Haryana | 2020-04-16 | Ancestral | -129.8 | 1.124 |
| EPI_ISL_454866 | hCoV-19/India/THSTI-BAL_914/2020 | Asia / India / Haryana | 2020-04-16 | Ancestral | -129.8 | 1.124 |
| EPI_ISL_454867 | hCoV-19/India/THSTI-BAL_918/2020 | Asia / India / Haryana | 2020-04-16 | Ancestral | -129.8 | 1.124 |
| EPI_ISL_455015 | hCoV-19/India/GBRC80/2020 | Asia / India / Gujarat / Rajkot | 2020-04-28 | Ancestral | -129.8 | 1.124 |
| EPI_ISL_455016 | hCoV-19/India/GBRC119/2020 | Asia / India / Gujarat / Rajkot | 2020-04-27 | Ancestral | -129.8 | 1.124 |
| EPI_ISL_455017 | hCoV-19/India/GBRC120/2020 | Asia / India / Gujarat / Vadodara | 2020-05-02 | D614G | -149.3 | 1.037 |

| | | | | | | |
|---|---|---|---|---|---|---|
| EPI_ISL_455018 | hCoV-19/India/GBRC121/2020 | Asia / India / Gujarat / Vadodara | 2020-05-02 | D614G:N148Y | -154 | 1.652 |
| EPI_ISL_455019 | hCoV-19/India/GBRC122/2020 | Asia / India / Gujarat / Vadodara | 2020-05-02 | D614G | -149.3 | 1.037 |
| EPI_ISL_455020 | hCoV-19/India/GBRC123/2020 | Asia / India / Gujarat / Vadodara | 2020-05-02 | D614G:S255F | -150.2 | 2.121 |
| EPI_ISL_455021 | hCoV-19/India/GBRC124/2020 | Asia / India / Gujarat / Vadodara | 2020-05-02 | D614G | -149.3 | 1.037 |
| EPI_ISL_455022 | hCoV-19/India/GBRC125/2020 | Asia / India / Gujarat / Vadodara | 2020-05-02 | D614G | -149.3 | 1.037 |
| EPI_ISL_455023 | hCoV-19/India/GBRC126/2020 | Asia / India / Gujarat / Vadodara | 2020-05-02 | D614G | -149.3 | 1.037 |
| EPI_ISL_455024 | hCoV-19/India/GBRC127/2020 | Asia / India / Gujarat / Vadodara | 2020-05-02 | D614G | -149.3 | 1.037 |
| EPI_ISL_455025 | hCoV-19/India/GBRC128/2020 | Asia / India / Gujarat / Vadodara | 2020-05-02 | D614G | -149.3 | 1.037 |
| EPI_ISL_455026 | hCoV-19/India/GBRC129/2020 | Asia / India / Gujarat / Vadodara | 2020-05-02 | D614G | -149.3 | 1.037 |
| EPI_ISL_455027 | hCoV-19/India/GBRC130/2020 | Asia / India / Gujarat / Vadodara | 2020-05-02 | D614G | -149.3 | 1.037 |
| EPI_ISL_455478 | hCoV-19/India/RMRC49/2020 | Asia / India / Odisha | 2020-04-09 | D614G | -149.3 | 1.037 |

| | | | | | | |
|---|---|---|---|---|---|---|
| EPI_ISL_455640 | hCoV-19/India/S4/2020 | Asia / India / West_Bengal / East Medinipur | 2020-03-27 | D614G:T941K | -152.5 | 3.522 |
| EPI_ISL_455641 | hCoV-19/India/S7/2020 | Asia/India / West_Bengal / Kolkata | 2020-03-31 | Ancestral | -129.8 | 1.124 |
| EPI_ISL_455643 | hCoV-19/India/S13/2020 | Asia / India / West_Bengal / North 24 Parganas | 2020-03-30 | D614G | -149.3 | 1.037 |
| EPI_ISL_455644 | hCoV-19/India/S14/2020 | Asia / India / West_Bengal / East Medinipur | 2020-04-02 | D614G | -149.3 | 1.037 |
| EPI_ISL_455645 | hCoV-19/India/S15/2020 | Asia / India / West_Bengal / Kolkata | 2020-04-03 | Ancestral | -129.8 | 1.124 |
| EPI_ISL_455646 | hCoV-19/India/S16/2020 | Asia / India / West_Bengal / Howrah | 2020-04-03 | D614G | -149.3 | 1.037 |
| EPI_ISL_455647 | hCoV-19/India/S17/2020 | Asia / India / West_Bengal / Kolkata | 2020-04-19 | D614G | -149.3 | 1.037 |
| EPI_ISL_455648 | hCoV-19/India/S18/2020 | Asia / India / West_Bengal / Kolkata | 2020-04-20 | D614G:G261S | -134.7 | 2.531 |
| EPI_ISL_455649 | hCoV-19/India/S19/2020 | Asia / India / West_Bengal / Kolkata | 2020-04-20 | Ancestral | -129.8 | 1.124 |
| EPI_ISL_455650 | hCoV-19/India/S20/2020 | Asia / India / West_Bengal / Kolkata | 2020-04-20 | D614G | -149.3 | 1.037 |

| | | | | | | |
|---|---|---|---|---|---|---|
| EPI_ISL_455651 | hCoV-19/India/S21/2020 | Asia / India / West_Bengal / Kolkata | 2020-04-21 | D614G | -149.3 | 1.037 |
| EPI_ISL_455652 | hCoV-19/India/S22/2020 | Asia / India / West_Bengal / South 24 Parganas | 2020-04-21 | D614G | -149.3 | 1.037 |
| EPI_ISL_455653 | hCoV-19/India/S23/2020 | Asia / India / West_Bengal / South 24 Parganas | 2020-04-21 | D614G | -149.3 | 1.037 |
| EPI_ISL_455654 | hCoV-19/India/S24/2020 | Asia/India / West_Bengal / Kolkata | 2020-04-21 | D614G | -149.3 | 1.037 |
| EPI_ISL_455655 | hCoV-19/India/S25/2020 | Asia / India / Rajasthan / Bikaner | 2020-04-30 | D614G:T323I | -159.1 | 9.390 |
| EPI_ISL_455656 | hCoV-19/India/S27/2020 | Asia / India / West_Bengal / South 24 Parganas | 2020-04-30 | Ancestral | -129.8 | 1.124 |
| EPI_ISL_455657 | hCoV-19/India/S28/2020 | Asia/India / West_Bengal / Kolkata | 2020-04-30 | D614G | -149.3 | 1.037 |
| EPI_ISL_455658 | hCoV-19/India/S29/2020 | Asia / India / West_Bengal / South 24 Parganas | 2020-04-30 | Ancestral | -129.8 | 1.124 |
| EPI_ISL_455659 | hCoV-19/India/S30/2020 | Asia / India / West_Bengal / South 24 Parganas | 2020-05-01 | D614G | -149.3 | 1.037 |

| | | | | | | |
|---|---|---|---|---|---|---|
| EPI_ISL_455660 | hCoV-19/India/S31/2020 | Asia / India / West_Bengal / South 24 Parganas | 2020-05-01 | D614G | -149.3 | 1.037 |
| EPI_ISL_455661 | hCoV-19/India/S32/2020 | Asia/India / West_Bengal / Kolkata | 2020-05-01 | D614G:H49Y | -152.4 | 2.654 |
| EPI_ISL_455662 | hCoV-19/India/S33/2020 | Asia / India / West_Bengal / Howrah | 2020-05-02 | D614G | -149.3 | 1.037 |
| EPI_ISL_455663 | hCoV-19/India/S34/2020 | Asia / India / West_Bengal / South 24 Parganas | 2020-05-02 | D614G:T323I | -159.1 | 9.390 |
| EPI_ISL_455664 | hCoV-19/India/S42/2020 | Asia/India / West_Bengal / Kolkata | 2020-05-03 | D614G | -149.3 | 1.037 |
| EPI_ISL_455665 | hCoV-19/India/S44/2020 | Asia/India / West_Bengal / Kolkata | 2020-05-01 | D614G | -149.3 | 1.037 |
| EPI_ISL_455666 | hCoV-19/India/S45/2020 | Asia/India / West_Bengal / Kolkata | 2020-05-01 | D614G:G261S | -134.7 | 2.531 |
| EPI_ISL_455667 | hCoV-19/India/S46/2020 | Asia / India / West_Bengal / South 24 Parganas | 2020-05-03 | D614G | -149.3 | 1.037 |
| EPI_ISL_455668 | hCoV-19/India/S48/2020 | Asia / India / West_Bengal / North 24 Parganas | 2020-05-05 | D614G:T323I | -159.1 | 9.390 |

| | | | | | | |
|---|---|---|---|---|---|---|
| EPI_ISL_455669 | hCoV-19/India/S49/2020 | Asia / India / West_Bengal / Kolkata | 2020-05-03 | D614G:G261S | -134.7 | 2.531 |
| EPI_ISL_455670 | hCoV-19/India/S50/2020 | Asia / India / West_Bengal / Kolkata | 2020-05-03 | D614G:G261S:G769V | -131.3 | 3.347 |
| EPI_ISL_455671 | hCoV-19/India/S51/2020 | Asia / India / West_Bengal / Kolkata | 2020-05-03 | Ancestral | -129.8 | 1.124 |
| EPI_ISL_455672 | hCoV-19/India/S52/2020 | Asia / India / West_Bengal / North 24 Parganas | 2020-05-03 | D614G | -149.3 | 1.037 |
| EPI_ISL_455673 | hCoV-19/India/S53/2020 | Asia / India / West_Bengal / South 24 Parganas | 2020-05-02 | D614G | -149.3 | 1.037 |
| EPI_ISL_455674 | hCoV-19/India/S55/2020 | Asia / India / West_Bengal / Kolkata | 2020-05-03 | D614G | -149.3 | 1.037 |
| EPI_ISL_455675 | hCoV-19/India/S56/2020 | Asia / India / West_Bengal / North 24 Parganas | 2020-05-03 | D614G | -149.3 | 1.037 |
| EPI_ISL_455676 | hCoV-19/India/S58/2020 | Asia / India / West_Bengal / Kolkata | 2020-05-03 | D614G | -149.3 | 1.037 |
| EPI_ISL_455678 | hCoV-19/India/S61/2020 | Asia / India / West_Bengal / North 24 Parganas | 2020-04-30 | D614G | -149.3 | 1.037 |

| | | | | | | |
|---|---|---|---|---|---|---|
| EPI_ISL_455679 | hCoV-19/India/S62/2020 | Asia / India / West_Bengal / East Medinipur | 2020-05-04 | D614G | -149.3 | 1.037 |
| EPI_ISL_455749 | hCoV-19/India/RMRC103/2020 | Asia / India / Odisha | 2020-05-06 | Ancestral | -129.8 | 1.124 |
| EPI_ISL_455750 | hCoV-19/India/RMRC104/2020 | Asia / India / Odisha | 2020-05-06 | Ancestral | -129.8 | 1.124 |
| EPI_ISL_455751 | hCoV-19/India/RMRC106/2020 | Asia / India / Odisha | 2020-05-06 | Ancestral | -129.8 | 1.124 |
| EPI_ISL_455752 | hCoV-19/India/RMRC108/2020 | Asia / India / Odisha | 2020-05-06 | Ancestral | -129.8 | 1.124 |
| EPI_ISL_455753 | hCoV-19/India/RMRC110/2020 | Asia / India / Odisha | 2020-05-06 | Ancestral | -129.8 | 1.124 |
| EPI_ISL_455754 | hCoV-19/India/RMRC112/2020 | Asia / India / Odisha | 2020-05-06 | D614G | -149.3 | 1.037 |
| EPI_ISL_455755 | hCoV-19/India/RMRC154/2020 | Asia / India / Odisha | 2020-05-07 | D614G | -149.3 | 1.037 |
| EPI_ISL_455756 | hCoV-19/India/RMRC155/2020 | Asia / India / Odisha | 2020-05-07 | D614G:L5F | NA | NA |
| EPI_ISL_455757 | hCoV-19/India/RMRC156/2020 | Asia / India / Odisha | 2020-05-07 | D614G | -149.3 | 1.037 |
| EPI_ISL_455758 | hCoV-19/India/RMRC157/2020 | Asia / India / Odisha | 2020-05-07 | Ancestral | -129.8 | 1.124 |
| EPI_ISL_455759 | hCoV-19/India/RMRC158/2020 | Asia / India / Odisha | 2020-05-07 | Ancestral | -129.8 | 1.124 |
| EPI_ISL_455760 | hCoV-19/India/RMRC159/2020 | Asia / India / Odisha | 2020-05-07 | Ancestral | -129.8 | 1.124 |

| | | | | | | |
|---|---|---|---|---|---|---|
| EPI_ISL_455761 | hCoV-19/India/RMRC160/2020 | Asia / India / Odisha | 2020-05-07 | A930T | -156.4 | 5.020 |
| EPI_ISL_455762 | hCoV-19/India/RMRC162/2020 | Asia / India / Odisha | 2020-05-07 | Ancestral | -129.8 | 1.124 |
| EPI_ISL_455763 | hCoV-19/India/RMRC163/2020 | Asia / India / Odisha | 2020-05-07 | Ancestral | -129.8 | 1.124 |
| EPI_ISL_455764 | hCoV-19/India/RMRC164/2020 | Asia / India / Odisha | 2020-05-07 | Ancestral | -129.8 | 1.124 |
| EPI_ISL_455765 | hCoV-19/India/RMRC165/2020 | Asia / India / Odisha | 2020-05-07 | D614G | -149.3 | 1.037 |
| EPI_ISL_455766 | hCoV-19/India/RMRC166/2020 | Asia / India / Odisha | 2020-05-07 | Ancestral | -129.8 | 1.124 |
| EPI_ISL_455767 | hCoV-19/India/RMRC167/2020 | Asia / India / Odisha | 2020-05-07 | Ancestral | -129.8 | 1.124 |
| EPI_ISL_455768 | hCoV-19/India/RMRC169/2020 | Asia / India / Odisha | 2020-05-07 | A831S:G857C | -164.4 | 2.425 |
| EPI_ISL_455769 | hCoV-19/India/RMRC170/2020 | Asia / India / Odisha | 2020-05-07 | Ancestral | -129.8 | 1.124 |
| EPI_ISL_455770 | hCoV-19/India/RMRC171/2020 | Asia / India / Odisha | 2020-05-07 | T761S | -137.6 | 5.838 |
| EPI_ISL_455771 | hCoV-19/India/RMRC2/2020 | Asia / India / Odisha | 2020-03-19 | D614G:H1101Y:K1191N | NA | NA |
| EPI_ISL_455773 | hCoV-19/India/RMRC22/2020 | Asia / India / Odisha | 2020-04-04 | D614G:I402L | -135.1 | 1.101 |
| EPI_ISL_455774 | hCoV-19/India/RMRC23/2020 | Asia / India / Odisha | 2020-04-04 | D614G | -149.3 | 1.037 |
| EPI_ISL_455775 | hCoV-19/India/RMRC24/2020 | Asia / India / Odisha | 2020-04-04 | D614G | -149.3 | 1.037 |

| | | | | | | |
|---|---|---|---|---|---|---|
| EPI_ISL_455776 | hCoV-19/India/RMRC27/2020 | Asia / India / Odisha | 2020-04-04 | D614G:L828P | -158 | 4.044 |
| EPI_ISL_455777 | hCoV-19/India/RMRC28/2020 | Asia / India / Odisha | 2020-04-04 | D614G | -149.3 | 1.037 |
| EPI_ISL_455778 | hCoV-19/India/RMRC30/2020 | Asia / India / Odisha | 2020-04-04 | D614G | -149.3 | 1.037 |
| EPI_ISL_455779 | hCoV-19/India/RMRC38/2020 | Asia / India / Odisha | 2020-04-04 | D614G | -149.3 | 1.037 |
| EPI_ISL_455780 | hCoV-19/India/RMRC46/2020 | Asia / India / Odisha | 2020-04-08 | D614G | -149.3 | 1.037 |
| EPI_ISL_455782 | hCoV-19/India/RMRC5/2020 | Asia / India / Odisha | 2020-04-02 | D614G | -149.3 | 1.037 |
| EPI_ISL_455783 | hCoV-19/India/RMRC6/2020 | Asia / India / Odisha | 2020-04-02 | D614G | -149.3 | 1.037 |
| EPI_ISL_455784 | hCoV-19/India/RMRC7/2020 | Asia / India / Odisha | 2020-04-02 | D614G | -149.3 | 1.037 |
| EPI_ISL_455786 | hCoV-19/India/RMRC90/2020 | Asia / India / Odisha | 2020-05-04 | D614G | -149.3 | 1.037 |
| EPI_ISL_455787 | hCoV-19/India/RMRC92/2020 | Asia / India / Odisha | 2020-05-04 | D614G | -149.3 | 1.037 |
| EPI_ISL_458030 | hCoV-19/India/CCMB_C12/2020 | Asia / India / Tamil Nadu | 2020-04-26 | D614G | -149.3 | 1.037 |
| EPI_ISL_458031 | hCoV-19/India/CCMB_C6/2020 | Asia / India / Tamil Nadu | 2020-04-26 | D614G | -149.3 | 1.037 |
| EPI_ISL_458032 | hCoV-19/India/CCMB_C8/2020 | Asia / India / Tamil Nadu | 2020-04-27 | D614G | -149.3 | 1.037 |
| EPI_ISL_458033 | hCoV-19/India/CCMB_C5/2020 | Asia / India / Tamil Nadu | 2020-04-29 | D614G | -149.3 | 1.037 |
| EPI_ISL_458034 | hCoV-19/India/CCMB_C11/2020 | Asia / India / Tamil Nadu | 2020-04-29 | D614G | -149.3 | 1.037 |

| | | | | | | |
|---|---|---|---|---|---|---|
| EPI_ISL_458035 | hCoV-19/India/CCMB_C13/2020 | Asia / India / Tamil Nadu | 2020-04-29 | K558N | -143.8 | 4.450 |
| EPI_ISL_458036 | hCoV-19/India/CCMB_C10/2020 | Asia / India / Tamil Nadu | 2020-04-29 | D614G | -149.3 | 1.037 |
| EPI_ISL_458037 | hCoV-19/India/CCMB_C7/2020 | Asia / India / Tamil Nadu | 2020-04-29 | D614G | -149.3 | 1.037 |
| EPI_ISL_458038 | hCoV-19/India/CCMB_C9/2020 | Asia / India / Tamil Nadu | 2020-04-29 | D614G | -149.3 | 1.037 |
| EPI_ISL_458039 | hCoV-19/India/CCMB_C16/2020 | Asia / India / Tamil Nadu | 2020-05-06 | D614G | -149.3 | 1.037 |
| EPI_ISL_458040 | hCoV-19/India/CCMB_C19/2020 | Asia / India / Tamil Nadu | 2020-05-06 | D614G | -149.3 | 1.037 |
| EPI_ISL_458041 | hCoV-19/India/CCMB_C15/2020 | Asia / India / Tamil Nadu | 2020-05-06 | D614G | -149.3 | 1.037 |
| EPI_ISL_458042 | hCoV-19/India/CCMB_C18/2020 | Asia / India / Tamil Nadu | 2020-05-06 | D614G | -149.3 | 1.037 |
| EPI_ISL_458043 | hCoV-19/India/CCMB_C14/2020 | Asia / India / Tamil Nadu | 2020-05-06 | D614G | -149.3 | 1.037 |
| EPI_ISL_458044 | hCoV-19/India/CCMB_C17/2020 | Asia / India / Tamil Nadu | 2020-05-06 | D614G | -149.3 | 1.037 |
| EPI_ISL_458045 | hCoV-19/India/CCMB_L1017/2020 | Asia / India / Telangana | 2020-05-11 | D614G:A520S:G1251V | NA | NA |
| EPI_ISL_458046 | hCoV-19/India/CCMB_L1021/2020 | Asia / India / Telangana | 2020-05-11 | D614G:A520S:G1251V | NA | NA |
| EPI_ISL_458047 | hCoV-19/India/CCMB_L1022/2020 | Asia / India / Telangana | 2020-05-11 | D614G:A520S:G1251V | NA | NA |

| | | | | | | |
|---|---|---|---|---|---|---|
| EPI_ISL_458048 | hCoV-19/India/CCMB_L1024/2020 | Asia / India / Telangana | 2020-05-11 | D614G:A520S:G1251V | NA | NA |
| EPI_ISL_458049 | hCoV-19/India/CCMB_L1025/2020 | Asia / India / Telangana | 2020-05-11 | D614G:A520S:G1251V | NA | NA |
| EPI_ISL_458050 | hCoV-19/India/CCMB_L1026/2020 | Asia / India / Telangana | 2020-05-11 | D614G:A520S:G1251V | NA | NA |
| EPI_ISL_458051 | hCoV-19/India/CCMB_L1062/2020 | Asia / India / Telangana | 2020-05-12 | D614G | -149.3 | 1.037 |
| EPI_ISL_458052 | hCoV-19/India/CCMB_L1063/2020 | Asia / India / Telangana | 2020-05-12 | D614G:V1068F | -158 | 4.785 |
| EPI_ISL_458053 | hCoV-19/India/CCMB_L1064/2020 | Asia / India / Telangana | 2020-05-12 | D614G | -149.3 | 1.037 |
| EPI_ISL_458054 | hCoV-19/India/CCMB_L1073/2020 | Asia / India / Telangana | 2020-05-13 | D614G | -149.3 | 1.037 |
| EPI_ISL_458055 | hCoV-19/India/CCMB_L1082/2020 | Asia / India / Telangana | 2020-05-13 | D614G:D1163Y | NA | NA |
| EPI_ISL_458056 | hCoV-19/India/CCMB_L1085/2020 | Asia / India / Telangana | 2020-05-13 | D614G | -149.3 | 1.037 |
| EPI_ISL_458057 | hCoV-19/India/CCMB_L1089/2020 | Asia / India / Telangana | 2020-05-13 | D614G | -149.3 | 1.037 |

| | | | | | | |
|---|---|---|---|---|---|---|
| EPI_ISL_458058 | hCoV-19/India/CCMB_L1090/2020 | Asia / India / Telangana | 2020-05-13 | D614G | -149.3 | 1.037 |
| EPI_ISL_458059 | hCoV-19/India/CCMB_L1100/2020 | Asia / India / Telangana | 2020-05-13 | D614G | -149.3 | 1.037 |
| EPI_ISL_458060 | hCoV-19/India/CCMB_L1103/2020 | Asia / India / Telangana | 2020-05-13 | D614G | -149.3 | 1.037 |
| EPI_ISL_458061 | hCoV-19/India/CCMB_L1108/2020 | Asia / India / Telangana | 2020-05-13 | D614G | -149.3 | 1.037 |
| EPI_ISL_458062 | hCoV-19/India/CCMB_L299/2020 | Asia / India / Telangana | 2020-04-20 | Ancestral | -129.8 | 1.124 |
| EPI_ISL_458063 | hCoV-19/India/CCMB_L301/2020 | Asia / India / Telangana | 2020-04-20 | Ancestral | -129.8 | 1.124 |
| EPI_ISL_458064 | hCoV-19/India/CCMB_L988/2020 | Asia / India / Telangana | 2020-05-11 | D614G | -149.3 | 1.037 |
| EPI_ISL_458065 | hCoV-19/India/CCMB_L989/2020 | Asia / India / Telangana | 2020-05-11 | D614G | -149.3 | 1.037 |
| EPI_ISL_458066 | hCoV-19/India/CCMB_O1/2020 | Asia / India / Telangana | 2020-04-13 | Ancestral | -129.8 | 1.124 |
| EPI_ISL_458067 | hCoV-19/India/CCMB_O2/2020 | Asia / India / Telangana | 2020-04-15 | Ancestral | -129.8 | 1.124 |
| EPI_ISL_458068 | hCoV-19/India/CCMB_O3/2020 | Asia / India / Telangana | 2020-04-14 | Ancestral | -129.8 | 1.124 |

| | | | | | | |
|---|---|---|---|---|---|---|
| EPI_ISL_458069 | hCoV-19/India/CCMB_O4/2020 | Asia / India / Telangana | 2020-04-03 | Ancestral | -129.8 | 1.124 |
| EPI_ISL_458070 | hCoV-19/India/CCMB_K272/2020 | Asia / India / Telangana | 2020-04-12 | Ancestral | -129.8 | 1.124 |
| EPI_ISL_458071 | hCoV-19/India/CCMB_J130/2020 | Asia / India / Telangana | 2020-04-01 | Ancestral | -129.8 | 1.124 |
| EPI_ISL_458072 | hCoV-19/India/CCMB_J812_P1/2020 | Asia / India / Telangana | 2020-04-08 | K77M | -84.8 | 0.194 |
| EPI_ISL_458073 | hCoV-19/India/CCMB_L276_P1/2020 | Asia / India / Telangana | 2020-04-20 | Ancestral | -129.8 | 1.124 |
| EPI_ISL_458074 | hCoV-19/India/CCMB_L301_P1/2020 | Asia / India / Telangana | 2020-04-20 | Ancestral | -129.8 | 1.124 |
| EPI_ISL_458075 | hCoV-19/India/CCMB_O2_P1/2020 | Asia / India / Telangana | 2020-04-15 | Ancestral | -129.8 | 1.124 |
| EPI_ISL_458076 | hCoV-19/India/CCMB_L302_P1/2020 | Asia / India / Telangana | 2020-04-20 | Ancestral | -129.8 | 1.124 |
| EPI_ISL_458077 | hCoV-19/India/CCMB_L040_P1/2020 | Asia / India / Telangana | 2020-04-18 | Ancestral | -129.8 | 1.124 |
| EPI_ISL_458080 | hCoV-19/India/CCMB_J300/2020 | Asia / India / Telangana | 2020-04-03 | Ancestral | -129.8 | 1.124 |
| EPI_ISL_458086 | hCoV-19/India/GBRC131/2020 | Asia / India / Gujarat / | 2020-05-24 | D614G | -149.3 | 1.037 |

| | | | | | | |
|---|---|---|---|---|---|---|
| | | Ahmedabad | | | | |
| EPI_ISL_458087 | hCoV-19/India/GBRC132/2020 | Asia / India / Gujarat / Ahmedabad | 2020-05-24 | D614G | -149.3 | 1.037 |
| EPI_ISL_458088 | hCoV-19/India/GBRC133/2020 | Asia / India / Gujarat / Ahmedabad | 2020-05-24 | D614G | -149.3 | 1.037 |
| EPI_ISL_458089 | hCoV-19/India/GBRC134/2020 | Asia / India / Gujarat / Ahmedabad | 2020-05-24 | D614G | -149.3 | 1.037 |
| EPI_ISL_458090 | hCoV-19/India/GBRC135a/2020 | Asia / India / Gujarat / Ahmedabad | 2020-05-24 | D614G:T572I | -145.6 | 8.737 |
| EPI_ISL_458091 | hCoV-19/India/GBRC135b/2020 | Asia / India / Gujarat / Ahmedabad | 2020-05-24 | D614G:T572I | -145.6 | 8.737 |
| EPI_ISL_458092 | hCoV-19/India/GBRC136a/2020 | Asia / India / Gujarat / Ahmedabad | 2020-05-24 | D614G | -149.3 | 1.037 |
| EPI_ISL_458093 | hCoV-19/India/GBRC136b/2020 | Asia / India / Gujarat / Ahmedabad | 2020-05-24 | D614G | -149.3 | 1.037 |
| EPI_ISL_458094 | hCoV-19/India/GBRC137a/2020 | Asia / India / Gujarat / Ahmedabad | 2020-05-24 | D614G | -149.3 | 1.037 |
| EPI_ISL_458095 | hCoV-19/India/GBRC137b/2020 | Asia / India / Gujarat / Ahmedabad | 2020-05-24 | D614G | -149.3 | 1.037 |
| EPI_ISL_458096 | hCoV-19/India/GBRC138/2020 | Asia / India / Gujarat / | 2020-05-24 | D614G | -149.3 | 1.037 |

| | | | | | | |
|---|---|---|---|---|---|---|
| | | Ahmedabad | | | | |
| EPI_ISL_458097 | hCoV-19/India/GBRC139a/2020 | Asia / India / Gujarat / Ahmedabad | 2020-05-24 | D614G | -149.3 | 1.037 |
| EPI_ISL_458098 | hCoV-19/India/GBRC139b/2020 | Asia / India / Gujarat / Ahmedabad | 2020-05-24 | D614G | -149.3 | 1.037 |
| EPI_ISL_458099 | hCoV-19/India/GBRC140/2020 | Asia / India / Gujarat / Ahmedabad | 2020-05-24 | D614G | -149.3 | 1.037 |
| EPI_ISL_458100 | hCoV-19/India/GBRC141a/2020 | Asia / India / Gujarat / Ahmedabad | 2020-05-24 | D614G:L54F | -137.6 | 1.061 |
| EPI_ISL_458101 | hCoV-19/India/GBRC141b/2020 | Asia / India / Gujarat / Ahmedabad | 2020-05-24 | D614G:L54F | -137.6 | 1.061 |
| EPI_ISL_458102 | hCoV-19/India/GBRC142/2020 | Asia / India / Gujarat / Ahmedabad | 2020-05-24 | D614G | -149.3 | 1.037 |
| EPI_ISL_458103 | hCoV-19/India/GBRC143/2020 | Asia / India / Gujarat / Bayad | 2020-05-05 | D614G:T572I:L5F | NA | NA |
| EPI_ISL_458104 | hCoV-19/India/GBRC144/2020 | Asia / India / Gujarat / Prantij | 2020-05-06 | D614G | -149.3 | 1.037 |
| EPI_ISL_458105 | hCoV-19/India/GBRC145/2020 | Asia / India / Gujarat / Prantij | 2020-05-06 | D614G | -149.3 | 1.037 |
| EPI_ISL_458106 | hCoV-19/India/GBRC146a/2020 | Asia / India / Gujarat / Himatnagar | 2020-05-16 | D614G | -149.3 | 1.037 |

| | | | | | | |
|---|---|---|---|---|---|---|
| EPI_ISL_458107 | hCoV-19/India/GBRC146b/2020 | Asia / India / Gujarat / Himatnagar | 2020-05-16 | D614G | -149.3 | 1.037 |
| EPI_ISL_458108 | hCoV-19/India/GBRC147/2020 | Asia / India / Gujarat / Khedbrahma | 2020-05-16 | D614G | -149.3 | 1.037 |
| EPI_ISL_458109 | hCoV-19/India/GBRC148a/2020 | Asia / India / Gujarat / Khedbrahma | 2020-05-17 | D614G | -149.3 | 1.037 |
| EPI_ISL_458110 | hCoV-19/India/GBRC148b/2020 | Asia / India / Gujarat / Khedbrahma | 2020-05-17 | Ancestral | -129.8 | 1.124 |
| EPI_ISL_458111 | hCoV-19/India/GBRC149/2020 | Asia / India / Gujarat / Himatnagar | 2020-05-17 | D614G | -149.3 | 1.037 |
| EPI_ISL_458112 | hCoV-19/India/GBRC150a/2020 | Asia / India / Gujarat / Himatnagar | 2020-05-18 | D614G:L54F | -137.6 | 1.061 |
| EPI_ISL_458113 | hCoV-19/India/GBRC150b/2020 | Asia / India / Gujarat / Himatnagar | 2020-05-18 | D614G:L54F | -137.6 | 1.061 |
| EPI_ISL_458298 | hCoV-19/India/CCMB_J714/2020 | Asia / India / Telangana | 2020-04-06 | Ancestral | -129.8 | 1.124 |
| EPI_ISL_459911 | hCoV-19/India/MaxCov0003_CSIR-IGIB/2020 | Asia / India / Delhi | 2020-05-09 | Ancestral | -129.8 | 1.124 |
| EPI_ISL_459915 | hCoV-19/India/MaxCov0013_CSIR-IGIB/2020 | Asia / India / Delhi | 2020-05-12 | Ancestral | -129.8 | 1.124 |

| | | | | | | |
|---|---|---|---|---|---|---|
| EPI_ISL_459916 | hCoV-19/India/MaxCov0014_CSIR-IGIB/2020 | Asia / India / Delhi | 2020-05-12 | Ancestral | -129.8 | 1.124 |
| EPI_ISL_459917 | hCoV-19/India/MaxCov0015_CSIR-IGIB/2020 | Asia / India / Delhi | 2020-05-08 | Ancestral | -129.8 | 1.124 |
| EPI_ISL_459918 | hCoV-19/India/MaxCov0016_CSIR-IGIB/2020 | Asia / India / Delhi | 2020-05-09 | Ancestral | -129.8 | 1.124 |
| EPI_ISL_459919 | hCoV-19/India/MaxCov0017_CSIR-IGIB/2020 | Asia / India / Delhi | 2020-05-21 | Ancestral | -129.8 | 1.124 |
| EPI_ISL_459920 | hCoV-19/India/MaxCov0018_CSIR-IGIB/2020 | Asia / India / Delhi | 2020-05-21 | Ancestral | -129.8 | 1.124 |
| EPI_ISL_459921 | hCoV-19/India/MaxCov0019_CSIR-IGIB/2020 | Asia / India / Delhi | 2020-05-08 | Ancestral | -129.8 | 1.124 |
| EPI_ISL_459922 | hCoV-19/India/MaxCov0020_CSIR-IGIB/2020 | Asia / India / Delhi | 2020-05-10 | Ancestral | -129.8 | 1.124 |
| EPI_ISL_459923 | hCoV-19/India/MaxCov0021_CSIR-IGIB/2020 | Asia / India / Delhi | 2020-05-08 | Ancestral | -129.8 | 1.124 |
| EPI_ISL_459925 | hCoV-19/India/MaxCov0023_CSIR-IGIB/2020 | Asia / India / Delhi | 2020-05-08 | Ancestral | -129.8 | 1.124 |

| | | | | | | |
|---|---|---|---|---|---|---|
| EPI_ISL_459926 | hCoV-19/India/MaxCov0024_CSIR-IGIB/2020 | Asia / India / Delhi | 2020 | Ancestral | -129.8 | 1.124 |
| EPI_ISL_459927 | hCoV-19/India/MaxCov0025_CSIR-IGIB/2020 | Asia / India / Delhi | 2020 | Ancestral | -129.8 | 1.124 |
| EPI_ISL_459928 | hCoV-19/India/MaxCov0026_CSIR-IGIB/2020 | Asia / India / Delhi | 2020 | Ancestral | -129.8 | 1.124 |
| EPI_ISL_459929 | hCoV-19/India/MaxCov0027_CSIR-IGIB/2020 | Asia / India / Delhi | 2020 | Ancestral | -129.8 | 1.124 |
| EPI_ISL_459930 | hCoV-19/India/MaxCov0028_CSIR-IGIB/2020 | Asia / India / Delhi | 2020 | Ancestral | -129.8 | 1.124 |
| EPI_ISL_459931 | hCoV-19/India/MaxCov0029_CSIR-IGIB/2020 | Asia / India / Delhi | 2020-05-09 | Ancestral | -129.8 | 1.124 |
| EPI_ISL_459932 | hCoV-19/India/MaxCov0030_CSIR-IGIB/2020 | Asia / India / Delhi | 2020-05-13 | Ancestral | -129.8 | 1.124 |
| EPI_ISL_459933 | hCoV-19/India/MaxCov0031_CSIR-IGIB/2020 | Asia / India / Delhi | 2020-05-05 | Ancestral | -129.8 | 1.124 |
| EPI_ISL_459934 | hCoV-19/India/MaxCov0032_CSIR-IGIB/2020 | Asia / India / Delhi | 2020-05-05 | Ancestral | -129.8 | 1.124 |

| | | | | | | |
|---|---|---|---|---|---|---|
| EPI_ISL_459935 | hCoV-19/India/MaxCov0033_CSIR-IGIB/2020 | Asia / India / Delhi | 2020-05-06 | Ancestral | -129.8 | 1.124 |
| EPI_ISL_459936 | hCoV-19/India/MaxCov0034_CSIR-IGIB/2020 | Asia / India / Delhi | 2020-05-10 | Ancestral | -129.8 | 1.124 |
| EPI_ISL_459937 | hCoV-19/India/MaxCov0035_CSIR-IGIB/2020 | Asia / India / Delhi | 2020-05-10 | Ancestral | -129.8 | 1.124 |
| EPI_ISL_459938 | hCoV-19/India/MaxCov0036_CSIR-IGIB/2020 | Asia / India / Delhi | 2020-05-11 | Ancestral | -129.8 | 1.124 |
| EPI_ISL_459939 | hCoV-19/India/MaxCov0037_CSIR-IGIB/2020 | Asia / India / Delhi | 2020-05-11 | Ancestral | -129.8 | 1.124 |
| EPI_ISL_459940 | hCoV-19/India/MaxCov0038_CSIR-IGIB/2020 | Asia / India / Delhi | 2020-05-10 | Ancestral | -129.8 | 1.124 |
| EPI_ISL_459941 | hCoV-19/India/MaxCov0039_CSIR-IGIB/2020 | Asia / India / Delhi | 2020-05-11 | Ancestral | -129.8 | 1.124 |
| EPI_ISL_459942 | hCoV-19/India/MaxCov0040_CSIR-IGIB/2020 | Asia / India / Delhi | 2020-04-27 | Ancestral | -129.8 | 1.124 |
| EPI_ISL_459943 | hCoV-19/India/MaxCov0041_CSIR-IGIB/2020 | Asia / India / Delhi | 2020-05-01 | Ancestral | -129.8 | 1.124 |

| | | | | | | |
|---|---|---|---|---|---|---|
| EPI_ISL_459944 | hCoV-19/India/MaxCov0042_CSIR-IGIB/2020 | Asia / India / Delhi | 2020-05-01 | Ancestral | -129.8 | 1.124 |
| EPI_ISL_459945 | hCoV-19/India/MaxCov0043_CSIR-IGIB/2020 | Asia / India / Delhi | 2020-04-29 | Ancestral | -129.8 | 1.124 |
| EPI_ISL_459946 | hCoV-19/India/MaxCov0044_CSIR-IGIB/2020 | Asia / India / Delhi | 2020-04-27 | Ancestral | -129.8 | 1.124 |
| EPI_ISL_459947 | hCoV-19/India/MaxCov0045_CSIR-IGIB/2020 | Asia / India / Delhi | 2020-04-28 | Ancestral | -129.8 | 1.124 |
| EPI_ISL_459948 | hCoV-19/India/MaxCov0046_CSIR-IGIB/2020 | Asia / India / Delhi | 2020-05-06 | Ancestral | -129.8 | 1.124 |
| EPI_ISL_459949 | hCoV-19/India/MaxCov0047_CSIR-IGIB/2020 | Asia / India / Delhi | 2020-04-27 | Ancestral | -129.8 | 1.124 |
| EPI_ISL_459950 | hCoV-19/India/MaxCov0048_CSIR-IGIB/2020 | Asia / India / Delhi | 2020-05-01 | Ancestral | -129.8 | 1.124 |
| EPI_ISL_459951 | hCoV-19/India/MaxCov0049_CSIR-IGIB/2020 | Asia / India / Delhi | 2020-05-01 | Ancestral | -129.8 | 1.124 |
| EPI_ISL_459952 | hCoV-19/India/MaxCov0050_CSIR-IGIB/2020 | Asia / India / Delhi | 2020-05-18 | Ancestral | -129.8 | 1.124 |

| | | | | | | |
|---|---|---|---|---|---|---|
| EPI_ISL_461478 | hCoV-19/India/GBRC151a/2020 | Asia / India / Gujarat / Vadodara | 2020-05-02 | D614G | -149.3 | 1.037 |
| EPI_ISL_461479 | hCoV-19/India/GBRC151b/2020 | Asia / India / Gujarat / Vadodara | 2020-05-02 | D614G | -149.3 | 1.037 |
| EPI_ISL_461480 | hCoV-19/India/GBRC152/2020 | Asia / India / Gujarat / Vadodara | 2020-05-02 | D614G:Q613H | -117.6 | 1.320 |
| EPI_ISL_461481 | hCoV-19/India/GBRC153a/2020 | Asia / India / Gujarat / Rajkot | 2020-04-27 | D614G | -149.3 | 1.037 |
| EPI_ISL_461482 | hCoV-19/India/GBRC153b/2020 | Asia / India / Gujarat / Rajkot | 2020-04-27 | Ancestral | -129.8 | 1.124 |
| EPI_ISL_461483 | hCoV-19/India/GBRC154/2020 | Asia / India / Gujarat / Ahmedabad | 2020-05-27 | D614G | -149.3 | 1.037 |
| EPI_ISL_461484 | hCoV-19/India/GBRC155/2020 | Asia / India / Gujarat / Ahmedabad | 2020-05-27 | D614G:T572I | -145.6 | 8.737 |
| EPI_ISL_461485 | hCoV-19/India/GBRC156/2020 | Asia / India / Gujarat / Ahmedabad | 2020-05-27 | D614G:E583D:D574Y:D1153Y | NA | NA |
| EPI_ISL_461486 | hCoV-19/India/GBRC157a/2020 | Asia / India / Gujarat / Ahmedabad | 2020-05-27 | D614G | -149.3 | 1.037 |
| EPI_ISL_461487 | hCoV-19/India/GBRC157b/2020 | Asia / India / Gujarat / Ahmedabad | 2020-05-27 | D614G | -149.3 | 1.037 |

| | | | | | | |
|---|---|---|---|---|---|---|
| EPI_ISL_461488 | hCoV-19/India/GBRC158/2020 | Asia / India / Gujarat / Ahmedabad | 2020-05-27 | D614G:P809S | -148.7 | 5.746 |
| EPI_ISL_461489 | hCoV-19/India/GBRC159a/2020 | Asia / India / Gujarat / Ahmedabad | 2020-05-27 | D614G | -149.3 | 1.037 |
| EPI_ISL_461490 | hCoV-19/India/GBRC159b/2020 | Asia / India / Gujarat / Ahmedabad | 2020-05-27 | D614G | -149.3 | 1.037 |
| EPI_ISL_461491 | hCoV-19/India/GBRC160a/2020 | Asia / India / Gujarat / Ahmedabad | 2020-05-27 | D614G | -149.3 | 1.037 |
| EPI_ISL_461492 | hCoV-19/India/GBRC160b/2020 | Asia / India / Gujarat / Ahmedabad | 2020-05-27 | D614G | -149.3 | 1.037 |
| EPI_ISL_461493 | hCoV-19/India/GBRC161a/2020 | Asia / India / Gujarat / Ahmedabad | 2020-05-27 | D614G | -149.3 | 1.037 |
| EPI_ISL_461494 | hCoV-19/India/GBRC161b/2020 | Asia / India / Gujarat / Ahmedabad | 2020-05-27 | D614G | -149.3 | 1.037 |
| EPI_ISL_461495 | hCoV-19/India/GBRC162a/2020 | Asia / India / Gujarat / Ahmedabad | 2020-05-27 | D614G:E583D:Q1201K | NA | NA |
| EPI_ISL_461496 | hCoV-19/India/GBRC162b/2020 | Asia / India / Gujarat / Ahmedabad | 2020-05-27 | D614G | -149.3 | 1.037 |
| EPI_ISL_461497 | hCoV-19/India/GBRC163a/2020 | Asia / India / Gujarat / Ahmedabad | 2020-05-27 | D614G | -149.3 | 1.037 |

| | | | | | | |
|---|---|---|---|---|---|---|
| EPI_ISL_461498 | hCoV-19/India/GBRC163b/2020 | Asia / India / Gujarat / Ahmedabad | 2020-05-27 | D614G | -149.3 | 1.037 |
| EPI_ISL_461499 | hCoV-19/India/GBRC164/2020 | Asia / India / Gujarat / Ahmedabad | 2020-05-27 | D614G | -149.3 | 1.037 |
| EPI_ISL_461500 | hCoV-19/India/GBRC165/2020 | Asia / India / Gujarat / Ahmedabad | 2020-05-27 | D614G:Q173H | -143.9 | 2.993 |
| EPI_ISL_461501 | hCoV-19/India/GBRC166a/2020 | Asia / India / Gujarat / Ahmedabad | 2020-05-27 | D614G | -149.3 | 1.037 |
| EPI_ISL_461502 | hCoV-19/India/GBRC166b/2020 | Asia / India / Gujarat / Ahmedabad | 2020-05-27 | D614G | -149.3 | 1.037 |
| EPI_ISL_461503 | hCoV-19/India/GBRC167a/2020 | Asia / India / Gujarat / Ahmedabad | 2020-05-27 | D614G | -149.3 | 1.037 |
| EPI_ISL_461504 | hCoV-19/India/GBRC167b/2020 | Asia / India / Gujarat / Ahmedabad | 2020-05-27 | D614G | -149.3 | 1.037 |
| EPI_ISL_461505 | hCoV-19/India/GBRC168a/2020 | Asia / India / Gujarat / Ahmedabad | 2020-05-27 | D614G | -149.3 | 1.037 |
| EPI_ISL_461506 | hCoV-19/India/GBRC168b/2020 | Asia / India / Gujarat / Ahmedabad | 2020-05-27 | D614G | -149.3 | 1.037 |

**S2 Table**.

Countrywide and state-wise frequency distribution matrix of circulating variants along with the docking score of S-R complex for each variant. The docking scores of the variants with mutations between amino acid positions 1-26 and 1148-1273 were not determined because of the unavailability of the modeling template [1, 2] and therefore mentioned as NA (not available). The variant mutation positions marked by asterisks denote convergent nature of these mutations, suggesting their independent accumulation multiple times along phylogenetically unlinked allelic backgrounds. The variants that were detected to be uniquely present in Indian population as compared to a total of 17529 worldwide genomic isolates available from the GISAID database till May 9, 2020.

| Mutation Type | | Unique in Indian population | Docking Score | Standard Error | State-wise Frequency | | | | | | | | | | | | | | | | | | | Total Frequency |
|---|---|---|---|---|---|---|---|---|---|---|---|---|---|---|---|---|---|---|---|---|---|---|---|---|
| | | | | | ANDHRA PRADESH | ASSAM | BIHAR | DELHI | GUJARAT | HARYANA | JAMMU | KARNATAKA | KERALA | LADAKH | MADHYA PRADESH | MAHARASHTRA | ODISHA | PUNJAB | RAJASTHAN | TAMIL NADU | TELANGANA | UTTAR PRADESH | WEST BENGAL | |
| **Ancestor 1 (Wuhan-Hu-1/2019)** | | | -129.8 | 1.124 | 1 | 2 | 4 | 57 | 16 | 10 | 1 | 8 | 0 | 7 | 3 | 31 | 15 | 0 | 4 | 16 | 65 | 3 | 10 | 253 |

| | | | | | | | | | | | | | | | | | | | | | | | | |
|---|---|---|---|---|---|---|---|---|---|---|---|---|---|---|---|---|---|---|---|---|---|---|---|---|
| **Ancestor 1 derived** | A243S | | -119.9 | 0.950 | 0 | 0 | 1 | 1 | 0 | 0 | 0 | 0 | 0 | 0 | 0 | 0 | 0 | 0 | 0 | 0 | 0 | 0 | 0 | 2 |
| | A831S:G857C | Y | -164.4 | 2.425 | 0 | 0 | 0 | 0 | 0 | 0 | 0 | 0 | 0 | 0 | 0 | 0 | 1 | 0 | 0 | 0 | 0 | 0 | 0 | 1 |
| | A930T | Y | -156.4 | 5.020 | 0 | 0 | 0 | 0 | 0 | 0 | 0 | 0 | 0 | 0 | 0 | 0 | 1 | 0 | 0 | 0 | 0 | 0 | 0 | 1 |
| | A930V | Y | -140.7 | 1.039 | 0 | 0 | 0 | 0 | 0 | 0 | 0 | 0 | 1 | 0 | 0 | 0 | 0 | 0 | 0 | 0 | 0 | 0 | 0 | 1 |
| | D290Y | Y | -119.1 | 1.389 | 0 | 0 | 0 | 0 | 0 | 0 | 0 | 0 | 0 | 0 | 0 | 1 | 0 | 0 | 0 | 0 | 0 | 0 | 0 | 1 |
| | E224G | Y | -95.9 | 0.857 | 0 | 0 | 0 | 1 | 0 | 0 | 0 | 0 | 0 | 0 | 0 | 0 | 0 | 0 | 0 | 0 | 0 | 0 | 0 | 1 |
| | E583D* | | -141 | 6.485 | 0 | 0 | 0 | 0 | 1 | 0 | 0 | 0 | 0 | 0 | 0 | 0 | 0 | 0 | 0 | 0 | 0 | 0 | 0 | 1 |
| | I1179N | Y | NA | NA | 0 | 0 | 0 | 0 | 0 | 0 | 0 | 0 | 0 | 0 | 0 | 1 | 0 | 0 | 0 | 0 | 0 | 0 | 0 | 1 |
| | K558N* | | -143.8 | 4.450 | 0 | 0 | 0 | 0 | 0 | 0 | 0 | 0 | 0 | 0 | 0 | 0 | 0 | 0 | 0 | 1 | 0 | 0 | 0 | 1 |
| | K77M | Y | -84.8 | 0.194 | 0 | 0 | 0 | 1 | 0 | 0 | 0 | 0 | 0 | 0 | 0 | 0 | 0 | 0 | 0 | 0 | 7 | 0 | 0 | 8 |
| | K77M:A771V | | -91.3 | 1.125 | 0 | 0 | 0 | 0 | 0 | 0 | 0 | 0 | 0 | 0 | 0 | 0 | 0 | 0 | 0 | 1 | 0 | 0 | 0 | 1 |
| | K77M:S1021F | Y | -128.5 | 0.354 | 0 | 0 | 0 | 0 | 0 | 0 | 0 | 0 | 0 | 0 | 0 | 0 | 0 | 0 | 0 | 0 | 1 | 0 | 0 | 1 |
| | K77M:S494P | Y | -76.3 | 4.382 | 0 | 0 | 1 | 0 | 0 | 0 | 0 | 0 | 0 | 0 | 0 | 0 | 0 | 0 | 0 | 0 | 0 | 0 | 0 | 1 |
| | L18F | | NA | NA | 0 | 0 | 0 | 1 | 0 | 0 | 0 | 0 | 0 | 0 | 0 | 0 | 0 | 0 | 0 | 0 | 0 | 0 | 0 | 1 |
| | L5F* | | NA | NA | 0 | 0 | 0 | 1 | 0 | 0 | 0 | 0 | 0 | 0 | 0 | 0 | 0 | 0 | 0 | 0 | 0 | 0 | 0 | 1 |
| | Q677H* | | -118.9 | 3.498 | 0 | 0 | 0 | 0 | 1 | 0 | 0 | 0 | 0 | 0 | 0 | 1 | 0 | 0 | 0 | 0 | 0 | 0 | 0 | 2 |
| | T1027 | | -136.7 | 1.131 | 0 | 0 | 0 | 0 | 0 | 0 | 0 | 0 | 0 | 0 | 0 | 0 | 0 | 0 | 1 | 0 | 0 | 0 | 0 | 1 |

| | | | | | | | | | | | | | | | | | | | | | | | | |
|---|---|---|---|---|---|---|---|---|---|---|---|---|---|---|---|---|---|---|---|---|---|---|---|---|
| | I | | | | | | | | | | | | | | | | | | | | | | | |
| | T274I | Y | -90.9 | 1.283 | 0 | 0 | 0 | 0 | 0 | 0 | 0 | 0 | 0 | 0 | 0 | 0 | 0 | 0 | 0 | 1 | 0 | 0 | 0 | 1 |
| | T602I | Y | -122.2 | 1.061 | 0 | 0 | 0 | 0 | 0 | 0 | 0 | 0 | 0 | 0 | 0 | 0 | 0 | 0 | 0 | 0 | 1 | 0 | 0 | 1 |
| | T761S | Y | -137.6 | 5.838 | 0 | 0 | 0 | 0 | 0 | 0 | 0 | 0 | 0 | 0 | 0 | 0 | 1 | 0 | 0 | 0 | 0 | 0 | 0 | 1 |
| | W258 del | | -116.7 | 1.637 | 0 | 0 | 0 | 0 | 0 | 0 | 0 | 0 | 0 | 0 | 0 | 1 | 0 | 0 | 0 | 0 | 0 | 0 | 0 | 1 |
| | Y145 del:R407I | | -105.2 | 7.245 | 0 | 0 | 0 | 0 | 0 | 0 | 0 | 0 | 1 | 0 | 0 | 0 | 0 | 0 | 0 | 0 | 0 | 0 | 0 | 1 |
| | Y28H | | -109.2 | 1.092 | 0 | 0 | 0 | 0 | 0 | 0 | 0 | 0 | 0 | 0 | 0 | 0 | 0 | 0 | 0 | 0 | 1 | 0 | 0 | 1 |
| **Ancestor 2 (D614G)** | | | -149.3 | 1.037 | 0 | 0 | 0 | 11 | 120 | 0 | 0 | 6 | 0 | 0 | 9 | 27 | 16 | 1 | 0 | 15 | 14 | 2 | 27 | 248 |
| **Ancestor 2 derived** | D614G:A520S:G1251V | Y | NA | NA | 0 | 0 | 0 | 0 | 0 | 0 | 0 | 0 | 0 | 0 | 0 | 0 | 0 | 0 | 0 | 0 | 6 | 0 | 0 | 6 |
| | D614G:A706S | | -136 | 4.137 | 0 | 0 | 0 | 0 | 1 | 0 | 0 | 0 | 0 | 0 | 0 | 0 | 0 | 0 | 0 | 0 | 0 | 0 | 0 | 1 |
| | D614G:A706S:C1243F* | Y | NA | NA | 0 | 0 | 0 | 0 | 1 | 0 | 0 | 0 | 0 | 0 | 0 | 0 | 0 | 0 | 0 | 0 | 0 | 0 | 0 | 1 |
| | D614G:A879S | | -146.6 | 21.300 | 0 | 0 | 0 | 0 | 1 | 0 | 0 | 0 | 0 | 0 | 0 | 0 | 0 | 0 | 0 | 0 | 0 | 0 | 0 | 1 |
| | D614G:A892V | Y | -143 | 3.897 | 0 | 0 | 0 | 0 | 1 | 0 | 0 | 0 | 0 | 0 | 0 | 0 | 0 | 0 | 0 | 0 | 0 | 0 | 0 | 1 |
| | D614G:C1243F* | Y | NA | NA | 0 | 0 | 0 | 0 | 1 | 0 | 0 | 0 | 0 | 0 | 1 | 0 | 0 | 0 | 0 | 0 | 0 | 0 | 0 | 2 |

| | | | | | | | | | | | | | | | | | | | | | | | | |
|---|---|---|---|---|---|---|---|---|---|---|---|---|---|---|---|---|---|---|---|---|---|---|---|---|
| | D614G:C1250F | Y | NA | NA | 0 | 0 | 0 | 0 | 0 | 0 | 0 | 1 | 0 | 0 | 0 | 0 | 0 | 0 | 0 | 0 | 0 | 0 | 0 | 1 |
| | D614G:D1163Y | | NA | NA | 0 | 0 | 0 | 0 | 0 | 0 | 0 | 0 | 0 | 0 | 0 | 0 | 0 | 0 | 0 | 0 | 1 | 0 | 0 | 1 |
| | D614G:D574H* | Y | -173.1 | 3.875 | 0 | 0 | 0 | 0 | 0 | 0 | 0 | 0 | 0 | 0 | 0 | 1 | 0 | 0 | 0 | 0 | 0 | 0 | 0 | 1 |
| | D614G:E156D | | -146.6 | 2.590 | 0 | 0 | 0 | 0 | 1 | 0 | 0 | 0 | 0 | 0 | 0 | 0 | 0 | 0 | 0 | 0 | 0 | 0 | 0 | 1 |
| | D614G:E484Q | Y | -150.4 | 0.714 | 0 | 0 | 0 | 0 | 0 | 0 | 0 | 0 | 0 | 0 | 0 | 1 | 0 | 0 | 0 | 0 | 0 | 0 | 0 | 1 |
| | D614G:E583D* | | -128.8 | 1.091 | 0 | 0 | 0 | 0 | 5 | 0 | 0 | 0 | 0 | 0 | 0 | 0 | 0 | 0 | 0 | 0 | 0 | 0 | 0 | 5 |
| | D614G:E583D*:D574Y*:D1153Y | Y | NA | NA | 0 | 0 | 0 | 0 | 1 | 0 | 0 | 0 | 0 | 0 | 0 | 0 | 0 | 0 | 0 | 0 | 0 | 0 | 0 | 1 |
| | D614G:E583D*:Q1201K | | NA | NA | 0 | 0 | 0 | 0 | 1 | 0 | 0 | 0 | 0 | 0 | 0 | 0 | 0 | 0 | 0 | 0 | 0 | 0 | 0 | 1 |
| | D614G:F2L | | NA | NA | 0 | 0 | 0 | 0 | 1 | 0 | 0 | 0 | 0 | 0 | 0 | 0 | 0 | 0 | 0 | 0 | 0 | 0 | 0 | 1 |
| | D614G:F797C | Y | -147.5 | 1.147 | 0 | 0 | 0 | 0 | 0 | 0 | 0 | 0 | 0 | 0 | 0 | 1 | 0 | 0 | 0 | 0 | 0 | 0 | 0 | 1 |

| | | | | | | | | | | | | | | | | | | | | | | | | |
|---|---|---|---|---|---|---|---|---|---|---|---|---|---|---|---|---|---|---|---|---|---|---|---|---|
| | D614G:G1124V | | -121.3 | 1.565 | 0 | 0 | 0 | 0 | 0 | 0 | 0 | 0 | 0 | 0 | 0 | 0 | 0 | 0 | 0 | 0 | 0 | 0 | 2 | 2 |
| | D614G:G261S | Y | -134.7 | 2.531 | 0 | 0 | 0 | 0 | 0 | 0 | 0 | 0 | 0 | 0 | 0 | 0 | 0 | 0 | 0 | 0 | 0 | 0 | 3 | 3 |
| | D614G:G261S:G769V | Y | -131.3 | 3.347 | 0 | 0 | 0 | 0 | 0 | 0 | 0 | 0 | 0 | 0 | 0 | 0 | 0 | 0 | 0 | 0 | 0 | 0 | 1 | 1 |
| | D614G:H1083Q:E583D* | Y | -152.4 | 3.817 | 0 | 0 | 0 | 0 | 1 | 0 | 0 | 0 | 0 | 0 | 0 | 0 | 0 | 0 | 0 | 0 | 0 | 0 | 0 | 1 |
| | D614G:H1083Q:R78M* | Y | -141.7 | 1.131 | 0 | 0 | 0 | 0 | 1 | 0 | 0 | 0 | 0 | 0 | 0 | 0 | 0 | 0 | 0 | 0 | 0 | 0 | 0 | 1 |
| | D614G:H1101Y:K1191N | | NA | NA | 0 | 0 | 0 | 0 | 0 | 0 | 0 | 0 | 0 | 0 | 0 | 0 | 1 | 0 | 0 | 0 | 0 | 0 | 0 | 1 |
| | D614G:H49Y | | -152.4 | 2.654 | 0 | 0 | 0 | 0 | 0 | 0 | 0 | 0 | 0 | 0 | 0 | 0 | 0 | 0 | 0 | 0 | 0 | 0 | 1 | 1 |
| | D614G:I1179N | Y | NA | NA | 0 | 0 | 0 | 0 | 0 | 0 | 0 | 0 | 0 | 0 | 0 | 1 | 0 | 0 | 0 | 0 | 0 | 0 | 0 | 1 |
| | D614G:I402L | Y | -135.1 | 1.101 | 0 | 0 | 0 | 0 | 0 | 0 | 0 | 0 | 0 | 0 | 0 | 0 | 1 | 0 | 0 | 0 | 0 | 0 | 0 | 1 |

| | | | | | | | | | | | | | | | | | | | | | | | | |
|---|---|---|---|---|---|---|---|---|---|---|---|---|---|---|---|---|---|---|---|---|---|---|---|---|
| | D614G:K1181R | Y | NA | NA | 0 | 0 | 0 | 0 | 1 | 0 | 0 | 0 | 0 | 0 | 0 | 0 | 0 | 0 | 0 | 0 | 0 | 0 | 0 | 1 |
| | D614G:L54F* | | -137.6 | 1.061 | 0 | 0 | 0 | 0 | 8 | 0 | 0 | 0 | 0 | 0 | 0 | 0 | 0 | 0 | 0 | 0 | 0 | 0 | 0 | 8 |
| | D614G:L54F*:E471Q | Y | -144.3 | 2.941 | 0 | 0 | 0 | 0 | 1 | 0 | 0 | 0 | 0 | 0 | 0 | 0 | 0 | 0 | 0 | 0 | 0 | 0 | 0 | 1 |
| | D614G:L5F* | | NA | NA | 0 | 0 | 0 | 0 | 0 | 0 | 0 | 0 | 0 | 0 | 0 | 0 | 1 | 0 | 0 | 0 | 0 | 0 | 0 | 1 |
| | D614G:L5F*:S162I | Y | NA | NA | 0 | 0 | 0 | 0 | 1 | 0 | 0 | 0 | 0 | 0 | 0 | 0 | 0 | 0 | 0 | 0 | 0 | 0 | 0 | 1 |
| | D614G:L828P | Y | -158 | 4.044 | 0 | 0 | 0 | 0 | 0 | 0 | 0 | 0 | 0 | 0 | 0 | 0 | 1 | 0 | 0 | 0 | 0 | 0 | 0 | 1 |
| | D614G:M177I | | -156.3 | 0.917 | 0 | 0 | 0 | 0 | 2 | 0 | 0 | 0 | 0 | 0 | 0 | 0 | 0 | 0 | 0 | 0 | 0 | 0 | 0 | 2 |
| | D614G:N148Y | Y | -154 | 1.652 | 0 | 0 | 0 | 0 | 1 | 0 | 0 | 0 | 0 | 0 | 0 | 0 | 0 | 0 | 0 | 0 | 0 | 0 | 0 | 1 |
| | D614G:P1263L | | NA | NA | 0 | 0 | 0 | 1 | 0 | 0 | 0 | 0 | 0 | 0 | 0 | 0 | 0 | 0 | 0 | 0 | 0 | 0 | 0 | 1 |
| | D614G:P809S | | -148.7 | 5.746 | 0 | 0 | 0 | 0 | 1 | 0 | 0 | 0 | 0 | 0 | 0 | 0 | 0 | 0 | 0 | 0 | 0 | 0 | 0 | 1 |
| | D614G:P82L | Y | -131.5 | 1.422 | 0 | 0 | 0 | 0 | 0 | 0 | 0 | 0 | 0 | 0 | 0 | 1 | 0 | 0 | 0 | 0 | 0 | 0 | 0 | 1 |

| | | | | | | | | | | | | | | | | | | | | | | | | |
|---|---|---|---|---|---|---|---|---|---|---|---|---|---|---|---|---|---|---|---|---|---|---|---|---|
| | D614G:Q173H | Y | -143.9 | 2.993 | 0 | 0 | 0 | 0 | 1 | 0 | 0 | 0 | 0 | 0 | 0 | 0 | 0 | 0 | 0 | 0 | 0 | 0 | 0 | 1 |
| | D614G:Q271R | Y | -109.5 | 0.477 | 0 | 0 | 0 | 0 | 2 | 0 | 0 | 0 | 0 | 0 | 0 | 0 | 0 | 0 | 0 | 0 | 0 | 0 | 0 | 2 |
| | D614G:Q613H | | -117.6 | 1.320 | 0 | 0 | 0 | 0 | 1 | 0 | 0 | 0 | 0 | 0 | 0 | 0 | 0 | 0 | 0 | 0 | 0 | 0 | 0 | 1 |
| | D614G:Q677H* | | -145.2 | 1.091 | 0 | 0 | 0 | 0 | 2 | 0 | 0 | 0 | 0 | 0 | 0 | 1 | 0 | 0 | 0 | 0 | 0 | 0 | 0 | 3 |
| | D614G:Q677H*:K558N* | Y | -133.8 | 2.711 | 0 | 0 | 0 | 0 | 0 | 0 | 0 | 0 | 0 | 0 | 0 | 1 | 0 | 0 | 0 | 0 | 0 | 0 | 0 | 1 |
| | D614G:R634S | Y | -144.5 | 1.814 | 0 | 0 | 0 | 0 | 0 | 0 | 0 | 0 | 0 | 0 | 0 | 1 | 0 | 0 | 0 | 0 | 0 | 0 | 0 | 1 |
| | D614G:R78M* | | -147.1 | 4.033 | 0 | 0 | 0 | 0 | 9 | 0 | 0 | 0 | 0 | 0 | 0 | 0 | 0 | 0 | 0 | 0 | 0 | 0 | 0 | 9 |
| | D614G:S13I | | NA | NA | 0 | 0 | 0 | 0 | 1 | 0 | 0 | 0 | 0 | 0 | 0 | 0 | 0 | 0 | 0 | 0 | 0 | 0 | 0 | 1 |
| | D614G:S255F | | -150.2 | 2.121 | 0 | 0 | 0 | 0 | 1 | 0 | 0 | 0 | 0 | 0 | 0 | 0 | 0 | 0 | 0 | 0 | 0 | 0 | 0 | 1 |
| | D614G:T22I | | NA | NA | 0 | 0 | 0 | 0 | 0 | 0 | 0 | 0 | 0 | 0 | 0 | 2 | 0 | 0 | 0 | 0 | 0 | 0 | 0 | 2 |
| | D614G:T22I:L7I | | NA | NA | 0 | 0 | 0 | 0 | 0 | 0 | 0 | 0 | 0 | 0 | 0 | 1 | 0 | 0 | 0 | 0 | 0 | 0 | 0 | 1 |

| | | | | | | | | | | | | | | | | | | | | | | | | |
|---|---|---|---|---|---|---|---|---|---|---|---|---|---|---|---|---|---|---|---|---|---|---|---|---|
| | D614G:T299I | Y | -120.9 | 1.636 | 0 | 0 | 0 | 1 | 0 | 0 | 0 | 0 | 0 | 0 | 0 | 0 | 0 | 0 | 0 | 0 | 0 | 0 | 0 | 1 |
| | D614G:T323I | | -159.1 | 9.390 | 0 | 0 | 0 | 0 | 0 | 0 | 0 | 0 | 0 | 0 | 0 | 0 | 0 | 0 | 1 | 0 | 0 | 0 | 2 | 3 |
| | D614G:T572I | | -145.6 | 8.737 | 0 | 0 | 0 | 0 | 7 | 0 | 0 | 0 | 0 | 0 | 0 | 0 | 0 | 0 | 0 | 0 | 0 | 0 | 0 | 7 |
| | D614G:T572I:L5F* | | NA | NA | 0 | 0 | 0 | 0 | 1 | 0 | 0 | 0 | 0 | 0 | 0 | 0 | 0 | 0 | 0 | 0 | 0 | 0 | 0 | 1 |
| | D614G:T723I | Y | -120 | 1.201 | 0 | 0 | 0 | 0 | 0 | 0 | 0 | 0 | 0 | 0 | 0 | 0 | 0 | 0 | 0 | 0 | 0 | 0 | 1 | 1 |
| | D614G:T827I | Y | -124.5 | 5.789 | 0 | 0 | 0 | 0 | 1 | 0 | 0 | 0 | 0 | 0 | 0 | 0 | 0 | 0 | 0 | 0 | 0 | 0 | 0 | 1 |
| | D614G:T941K | Y | -152.5 | 3.522 | 0 | 0 | 0 | 0 | 0 | 0 | 0 | 0 | 0 | 0 | 0 | 0 | 0 | 0 | 0 | 0 | 0 | 0 | 1 | 1 |
| | D614G:V1068F | Y | -158 | 4.785 | 0 | 0 | 0 | 0 | 0 | 0 | 0 | 0 | 0 | 0 | 0 | 0 | 0 | 0 | 0 | 0 | 1 | 0 | 0 | 1 |
| | D614G:V1104L | Y | -152.1 | 1.600 | 0 | 0 | 0 | 0 | 1 | 0 | 0 | 0 | 0 | 0 | 0 | 0 | 0 | 0 | 0 | 0 | 0 | 0 | 0 | 1 |

**S3 Table.**

State-wise distribution of the type and frequency of spike protein variants, along with the calculated diversity based on Simpson's index [3].

| State/Union Territories | No. of sequenced isolates | No. of variant types | Frequency of variants | Diversity (Simpson index) | Standard Error |
|---|---|---|---|---|---|
| Andhra Pradesh | 1 | 1 | 1 | not calculable | |
| Assam | 2 | 1 | 2 | 1 | 0 |
| Bihar | 6 | 3 | 4, 1, 1 | 0.5 | 0.19245 |
| Delhi | 75 | 9 | 57, 11, 1, 1, 1, 1, 1, 1, 1 | 0.60036 | 0.06602 |
| Gujarat | 196 | 34 | 16, 120, 1, 1, 1, 1, 1, 1, 5, 1, 1, 1, 1, 1, 1, 8, 1, 1, 2, 1, 1, 1, 2, 1, 2, 9, 1, 1, 7, 1, 1, 1, 1, 1 | 0.38817 | 0.0403 |
| Haryana | 10 | 1 | 10 | 1 | 0 |
| Jammu and Kashmir | 1 | 1 | 1 | not calculable | |
| Karnataka | 15 | 3 | 8, 6, 1 | 0.44889 | 0.06218 |
| Kerala | 2 | 2 | 1, 1 | not calculable | |
| Ladakh | 7 | 1 | 7 | 1 | 0 |
| Madhya Pradesh | 13 | 3 | 3, 9, 1 | 0.53846 | 0.12964 |
| Maharashtra | 73 | 16 | 31, 27, 1, 1, 1, 1, 1, 1, 1, 1, 1, 1, 1, 1, 1, 2 | 0.32032 | 0.03673 |
| Odisha | 38 | 9 | 15, 16, 1, 1, 1, 1, 1, 1, 1 | 0.33795 | 0.0482 |
| Punjab | 1 | 1 | 1 | not calculable | |
| Rajasthan | 6 | 3 | 4, 1, 1 | 0.5 | 0.19245 |
| Tamil Nadu | 34 | 5 | 16, 15, 1, 1, 1 | 0.41869 | 0.04181 |
| Telengana | 97 | 9 | 65, 14, 6, 1, 1, 7, 1, 1, 1 | 0.47944 | 0.05549 |
| Uttar Pradesh | 5 | 2 | 3, 2 | 0.52 | 0.08764 |
| West Bengal | 48 | 9 | 10, 27, 2, 3, 1, 1, 2, 1, 1 | 0.36892 | 0.06541 |

**S1 Dataset. Positional information of the spike protein mutations in the docked S-R complex.**

There are numerous non-synonymous mutations found over the entire range of Spike protein of the beta-coronavirus emerged worldwide till date. These mutations are not restricted at the receptor-binding domain (RBD) S-protein, hence may not modulate binding to the host directly, but allosterically, which could not be predicted from the docked complex alone.

We have modeled the spike protein and then docked each of 58 variants S-R complex within the available range of Swiss-model template [residue range: 27-1147] [1, 2].

The S1 subunit (residue range: 14-685) of spike protein has N-terminal (NTD, residue range:14-305), receptor binding domains (RBD, residue range: 319-541) and receptor binding motif (RBM, residue range: 437-508) while S2 subunit (residue rang: 686-1195) comprises of fusion peptide (FP), heptad repeat 1 (HR1) and heptad repeat 2 (HR2, residue range:1163-1213). Among our analyzed variants, ancestor1 contains 12 single point mutation variants positioned at S1 subunit, while 4 single point mutants were spotted on S2 subunit. Among the five analyzed double mutant/deletion-mutation, two were entirely located at S1 subunit, two were spotted on both S1 and S2 subunit, and one was solely located at S2 subunit. Ancestor 2 has twelve singly mutated analyzed variants located at S2 subunit and twenty-one analyzed variants were spotted on S1 subunit in D614G background. Two double-mutant variants in D614G background were positioned at S1 subunit and three were found in S2 subunit.

A closer look at the mutations was illustrated below.

**Ancestor 1 (Wuhan-Hu-1/2019)**

The docked structure of Ancestor 1 to the hACE2 was successfully reproduced the native salt bridges and hydrogen bonds reported in crystal structure [4].

**Y28H**

This mutation was found in a β-strand of the N-terminal domain of S1 subunit and was far away from the binding site. No significant structural change has been observed from the docked complex upon mutation.

**K77M**

This mutation was in N-terminal domain of S1 subunit, not in the binding domain, but far away from the binding site. The mutation has lead to a small helix (Thr76-8Asn81) formation which was absent in Ancestor 1.

**K77M:S494P**

S494P mutation has emerged in K77M background. Pro494 is located at β6 strand at RBD. As mentioned in case of K77M, this double mutation also introduced the similar small helix of the range Thr76-8Asn81.

**K77M:A771V**

The A771V mutation has emerged in K77M background. K77M is in the N-terminal domain of S1 subunit, while A771V was spotted in S2 subunit. The K77M mutation was in one of the loop of the N-terminal domain, while A771V is far away from the binding site. A771V mutation did not modify secondary structure, but K77M, being in the loop, altered the loop flexibility. The loop where the K77M mutation located was recoiled to form a small helix of (Thr76-8Asn81).

**K77M:1021F**

The S1021F mutation has emerged in K77M background. This mutation was found in the region of S2 subunit, connecting HR1 and HR2 domains. There was no visible conformational change nearby the mutational site. However both mutations are away from the binding site.

**Y145del:R407I**

The mutation comprised of one amino acid deletion at Tyr145 followed by mutation of arginine by isoleucine at 407, which was one of the binding site residues. The deletion was in S1 subunit, away from the binding site, in a loop, while mutation was at the RBD.

**E224G**

Mutation was positioned at the N-terminal domain of S1 subunit. Glutamate mutation by small and neutral glycine, far away from the binding site, did not show any significant secondary structural alternation.

**A243S**

The mutation was located at the N-terminal domain of S1 subunit, but replacement of alanine by serine causes two-residue shrink of a β-strand located (Thr236-His245 in ancestor-1 shrinks to Thr236-Ser243).

**W258del**

This mutation was located at the N-terminal domain of S1 subunit, in a flexible loop. There was no structural change observed in the docked complex except the fluctuation of the loop.

**T274I**

The mutation was located at the N-terminal domain of S1 subunit in a β-strand. There was two-residue elongation of the β-strand (Leu270-Lys278) upon mutation.

**D290Y**

The mutation was located at the N-terminal domain of S1 subunit, in a β-strand, and induced strong interactions at the binding site. There was no observed secondary structural change in the docked complex.

**K558N**

The mutation was located at S1 subunit, in a loop. No structural change has been observed in the docked complex.

**E583D**

The mutation was spotted on the S1 subunit in a β-strand. No significant change in secondary structure was observed in docked structure upon mutation.

**T602I**

This mutation was position at the S1 subunit in a connecting loop of two β-strands. There was no structural change observed in the docked complex.

**D614G**

Receptor binding domain of S1 subunit contained this mutation in a connecting loop. The docked complex highly resembled Ancestor-1.

**Q677H**

This mutation was position at the S1 subunit. A short helix (Tyr674-Ans679) formation was observed upon mutation alike the native Ancestor-1.

**T761S**

The mutation was located in a well-packed helical region of S2 subunit. No visible structural change has been observed.

**A831S:G857C**

Any significant change in secondary structure was not observed in this case. Both mutations are located at the S2 subunit in the region connected to fusion peptide and HR1 domain.

**A930V**

This mutation was located at HR1 domain of S2 subunit in a close-packed helix. No visible change in secondary structure has been observed.

**A930T**

This mutation was located at HR1 domain of S2 subunit in a close-packed helix as well. No visible secondary structural change has been found.

**T1027I**

This mutation was located at the connecting region of HR1 and HR1 domain of S2 subunit, in a close-packed helix. There was no structural change observed in the docked complex.

**Ancestor-2 (D614G variants)**

**D614G:H49Y**

The mutation was located at N-terminal domain of S1 subunit, in a β-strand. No structural change has been observed in the docked complex.

**D614G:L54F**

The mutation was located in the β-strand of N-terminal domain of S1 subunit. No change in secondary structure has been observed in the docked complex.

**D614G:L54F:E471Q**

This was a double mutation in D614G background. L54F was located at N-terminal domain and E471Q was positioned at binding site of S1 subunit. Both were found at the fluctuating loops as observed in the docked complex.

**D614G:R78M**

The mutation was located at N-terminal domain of S1 subunit in a flexible loop as observed in the docked structure.

**D614G:P82L**

The mutation was positioned at N-terminal domain of S1 subunit in a β-strand. No structural change was observed in the docked complex.

**N148Y**

The mutation was spotted at one of the loops of N-terminal domain of S1 subunit. No change in secondary structure was observed in the docked complex, except loop fluctuation.

**D614G:E156D**

The mutation was located at N-terminal domain of S1 subunit in a fluctuating loop.

**D614G:Q173H**

The mutation was located at N-terminal domain of S1 subunit and positioned in a loop, thus fluctuations were observed in the docked complex.

**D614G:M177I**

The mutation was located at N-terminal domain of S1 subunit in a flexible loop as observed from docked structure.

**D614GL:S255F**

The mutation was located at the N-terminal domain of S1 subunit. The mutation was positioned at the connecting loop of the β-strands, thus fluctuation has been observed in the docked complex.

**D614G:G261S**

The mutation was located at the N-terminal domain of S1 subunit. It was found to connect loop of the two β-strands, thus fluctuation has been observed in the docked complex.

**D614G:G261S:G769V**

This was a double mutation. G261S was located at N-terminal domain of S1 subunit and the fusion peptide of S2 subunit harbored G769V. The double mutation was in the flexible loop as it was observed in the docked complex.

**D614G:Q271R**

The mutation was located at N-terminal domain of S1 subunit, in a long β-strand. No structural change has been observed in the docked complex.

**D614G:T299I**

The mutation was found in a helix of the N-terminal domain of S1 subunit. No structural change has been observed in the docked complex.

**D614G_T323I**

The mutation was spotted at the receptor-binding domain of S1 subunit, connecting two β-strands. No structural change has been observed in the docked complex.

**D614G_I402L**

The mutation was located at the binding site in a fluctuating loop of the S1 subunit. No structural change has been observed in the docked complex.

**D614G:E484Q**

This mutation was located at receptor binding motif of S1 subunit in a fluctuating loop. No other structural change has been observed.

**D614G:T572I**

The mutation was found in the receptor binding domain of S1 subunit in a β-hairpin. No structural alteration has been observed in docked complex.

**D614G:E583D**

This mutation was located at S1 subunit, in a β-strand. The region contains flexible loops, as observed from the docked complex.

**D614G:D574H**

The mutation was found in a loop of the S1 subunit. No structural change has been found except nearby loop fluctuation.

**D614G:Q613H**

β-strand of S1 domain harbored this mutation. No significant change at the secondary structural level has been observed in the docked complex.

**D614G:R634S**

The mutation was spotted at the S1 subunit. It was located at the loop, thus fluctuation was observed in the docked structure.

**D614G_Q677H**

The mutation was located at S1 domain and far away from the binding site. This mutation caused a shrink of the β-strand compared to the same of the Ancestor-2.

**D614G:Q677H:K558N**

This was a double mutation, located at N-terminal domain of S1 subunit and fusion peptide of S2 subunit. Both mutations were positioned in fluctuating loops.

**D614G:A706S**

The mutation was located at S2 subunit, connecting S1 subunit and fusion peptide.

**D614G:T723I**

The mutation was located at S2 subunit in a well-packed β-strand. No structural change has been observed in the docked complex.

**D614G:F797C**

The mutation was located at S2 domain. The location of the mutation was far away from the binding site and the mutation does not cause any alteration in the secondary structure of the docked complex.

**D614G:P809S**

The mutation was located at the S2 subunit. Upon mutation, as proline is replace by serine, a β-strand formation has been observed in the docked complex.

**D614G:T827I**

The mutation was located at S2 subunit in a connecting loop. No significant change was observed in the docked complex except loop fluctuation.

**D614G:L828P**

This mutation was located at the S2 subunit. No significant change at the secondary structural level has been observed.

**D614G:A879S**

The mutation was located at S2 domain in a helical region. No structural change has been observed in the docked complex.

**D614G:A892V**

The mutation was located at S2 domain. There was no secondary structural change observed upon docking.

**D614G:T941K**

The mutation was positioned at HR1 domain of S2 subunit, in a closely packed helical region. There was no secondary structural change observed upon docking.

**D614G:V1068F**

The mutation was located at S2 subunit in a loop connecting HR1 and HR2 domains. No secondary structural change was observed except loop fluctuation upon docking.

**D614G:H1083Q:R78M**

This was a double mutation in D614G background. H1083Q was located at S2 subunit in the region connecting HR1 and HR2 domain, while R78M lies on N-terminal domain of S1 subunit. Upon R78M mutation, loop containing the mutation has been converted to the β-strand. H1083Q however was positioned in a relatively stable spot.

**D614G:V1104L**

The mutation was located at S2 domain. There was no secondary structural change observed upon docking.

**D614G:G1124V**

The mutation was located at S2 subunit in a β-strand. No structural change has been observed in the docked complex.
